# CRISPR-Cas Controls Cryptic Prophages

**DOI:** 10.1101/2021.07.28.454074

**Authors:** Sooyeon Song, Ekaterina Semenova, Konstantin Severinov, Laura Fernández-García, Michael J. Benedik, Toshinari Maeda, Thomas K. Wood

## Abstract

The bacterial archetypal adaptive immune system, CRISPR-Cas, is thought to be repressed in the best-studied bacterium, *Escherichia coli* K-12. We show here that the *E. coli* CRISPR-Cas system is active and serves to inhibit its nine defective (i.e., cryptic) prophages. Specifically, compared to the wild-type strain, reducing the amounts of specific interfering RNAs (crRNA) decreases growth by 40%, increases cell death by 700%, and prevents persister cell resuscitation. Similar results were obtained by inactivating CRISPR-Cas by deleting the entire 13 spacer region (CRISPR array); hence, CRISPR-Cas serves to inhibit the remaining deleterious effects of these cryptic prophages, most likely through CRISPR array-derived crRNA binding to cryptic prophage mRNA, rather than through cleavage of cryptic prophage DNA; i.e., self-targeting. Consistently, four of the 13 *E. coli* spacers contain complementary regions to the mRNA sequences of seven cryptic prophages, and inactivation of CRISPR-Cas increases the level of mRNA for lysis protein YdfD of cryptic prophage Qin and lysis protein RzoD of cryptic prophage DLP-12. Also, lysis is clearly seen via transmission electron microscopy when the whole CRISPR-Cas array is deleted, and eliminating spacer #12, which encodes crRNA with complementary regions for DLP-12 (including *rzoD*), Rac, Qin (including *ydfD*), and CP4-57 cryptic prophages, also results in growth inhibition and cell lysis. Therefore, we report the novel results that (i) CRISPR-Cas is active in *E. coli* and (ii) CRISPR-Cas is used to tame cryptic prophages, likely through RNAi; i.e., unlike with active lysogens, active CRISPR-Cas and cryptic prophages may stably co-exist.

## 1. INTRODUCTION

Along with restriction/modification (Vasu and Nagaraja, 2013) and toxin/antitoxin (TA) systems (Pecota and Wood, 1996), prokaryotes utilize clustered, regularly-interspaced, short palindromic repeats (CRISPR) and CRISPR-associated (Cas) (Makarova et al., 2020) proteins to combat phages. These systems are interrelated in that some Cas proteins and TA systems have a common ancestor; for example, *Sulfolobus solfataricus* Cas2 is structurally similar to antitoxin GhoS (an RNase) of the *Escherichia coli* GhoT/GhoS TA system (Wang et al., 2012). In addition, in a manner similar to our discovery (Pecota and Wood, 1996) that toxins of host TA systems inhibit phage by degrading mRNA when the phage shuts down transcription (e.g., Hok/Sok inhibits T4 phage), some Cas proteins induce host dormancy rather than degrading phage DNA to inhibit phage propagation (Meeske et al., 2019). Also, TA systems have been found to stabilize CRISPR-Cas systems by making them addictive to the host (Li et al., 2021).

Although CRISPR-Cas systems exclude both external lytic and temperate (lysogenic) phages (Goldberg et al., 2018), CRISPR-Cas systems of lysogens that target their own integrated prophages decrease long-term fitness and either the cell dies or the prophage is lost (Edgar and Qimron, 2010; HYPERLINK \l "bookmark11" Goldberg *et al*., 2018). Also, the class I-E (Makarova *et al*., 2020) CRISPR–Cas system of *E. coli* is not related to immunity for external phages (Touchon et al., 2011) and appears to be inactive in the wild-type strain at standard laboratory conditions (Shmakov et al., 2017), due to repression by H-NS, although it is functional when induced (Pougach et al., 2010). To date, the relationship of CRISPR-Cas to cryptic prophages; i.e., those phage remnants that are unable to form lytic particles, has not been investigated.

Up to 50% of bacterial genomes may contain stably-integrated phage DNA (Howard-Varona et al., 2017), and for *E. coli*, we discovered that its nine cryptic prophages are not extraneous DNA but instead encode genes for proteins that increase resistance to sublethal concentrations of quinolone and β-lactam antibiotics as well as protect the cell from osmotic, oxidative, and acid stresses (by deleting 166 kb) (Wang et al., 2010). Although these cryptic prophages do not enhance the formation of persister cells, a subpopulation of cells that weather extreme stress by entering into a dormant state (Wood and Song, 2020), these phage remnants facilitate the resuscitation of persisters via nutrient sensing (Song et al., 2021). Therefore, the bacterial cell can co-opt the genome of its former parasite to both combat stress (Wang *et al*., 2010) as well as to revive from dormancy (Song *et al*., 2021).

Given these active roles of cryptic prophages in the stress response (Wang *et al*., 2010) and persister cell resuscitation (Song *et al*., 2021), we hypothesized here that the native *E. coli* CRISPR-Cas system plays an active role in the regulation of cryptic prophages. We find that the CRISPR-Cas system is required for inhibiting the expression of deleterious cryptic prophage genes, since, if CRISPR-Cas is inactivated by preventing crRNA production, cells die due to activation of the cryptic prophage lysis proteins YdfD of Qin and RzoD from DLP-12. Hence, we discovered CRISPR-Cas is active in *E. coli* and serves to regulate its former phage foe.

## 2. RESULTS

### Deletions of CRISPR-Cas components reduce growth

We assayed the importance of CRISPR-Cas in *E. coli* K-12 by testing the individual deletions of the CRISPR-Cas genes on growth in M9 glucose medium (*cas1, cas2, cas3, casA, casB, casC, casD*, and *casE*) (**Fig. S1**) and found only the *cas2* deletion, with its kanamycin cassette insertion, had an effect: *cas2* causes 40% slower growth in rich medium compared to the wild-type strain (specific growth rate of 0.79 ± 0.21/h vs. 1.3 ± 0.11/h, respectively). Similarly, in minimal glucose (0.4 wt%) medium, deletion of *cas2* also reduces growth by 33% (0.42 ± 0.02/h vs. 0.62 ± 0.02/h, respectively) and reduced the yield in stationary phase (**Fig. S2A**). We hypothesized that the *cas2* deletion/kanamycin insertion caused a polar effect by reducing spacer production given it is directly upstream of the spacer region (Pougach *et al*., 2010) (**Fig. S3**). Fittingly, inactivation of CRISPR-Cas by eliminating the CRISPR arrays in the isogenic host BW25113 (henceforth Δspacer) also reduced growth by 32% in rich medium and eliminating kanamycin resistance in the *cas2* deletion strain to prevent the polar mutation affecting the spacer region restored nearly wild-type growth (**Fig. S2A**) as well as reduced toxicity (**Fig. S2B**) relative to the *cas2* Ωkan^R^ strain. Therefore, the *E. coli* CRISPR-Cas represses some processes that inhibit growth via crRNA, and by this criterion, is active, providing a clear advantage to the host.

### CRISPR-Cas increases single-cell resuscitation

Since deletions of CRISPR-Cas components decrease growth, we tested for their effect on persister cell resuscitation using single-cell microscopy. Persister cell resuscitation is germane in that the dormant cells are highly stressed and have limited resources for their revival via activation of hibernating ribosomes (Kim et al., 2018; Song and Wood, 2020; HYPERLINK \l "bookmark44" Yamasaki et al., 2020); for example, we have shown that inhibiting ATP synthesis leads to a 5,000-fold increase in persister cell formation (Kwan et al., 2013). Since we discovered facile means for converting the whole population of cells into persister cells (Kim *et al*., 2018; HYPERLINK \l "bookmark18" Kwan *et al*., 2013; HYPERLINK \l "bookmark44" Yamasaki *et al*., 2020) that has been used by at least 17 independent labs to date with various bacterial species (Song and Wood, 2021), these stressed cells are an excellent model for testing the effects of CRISPR-Cas on *E. coli* physiology.

Here, we found that deletion of *cas2* reduces persister cell resuscitation by 31-fold (**Fig. 1A, Table S1**). Persister cell resuscitation was similarly affected (15-fold reduction) in the Δspacer strain (**Fig. 1A, Table S1**). Since the *cas2* mutant grows more slowly than the wild-type in rich medium, as a control, we tested an *E. coli* mutant that grows 22% slower than the wild-type, *ssrA*, to confirm that the slower growth does not affect persister resuscitation and found that the *ssrA* mutant resuscitates at nearly the identical rate as the wild-type strain (**Fig. S4**). Hence, CRISPR-Cas is active in *E. coli* and plays key roles in its growth and recovery from extreme stress.

**Figure 1.**
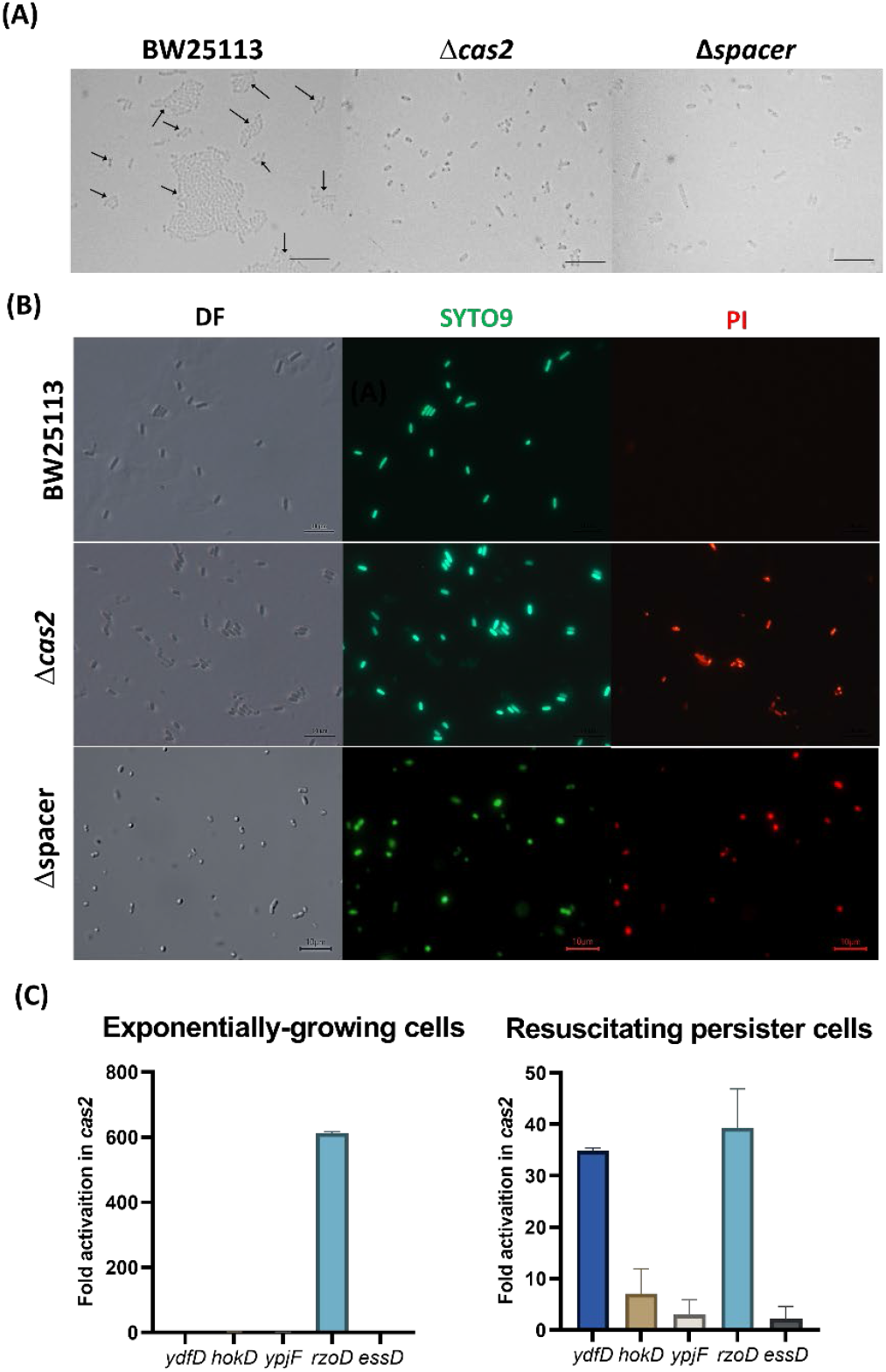
Inactivating CRISPR-Cas eliminates persister cell resuscitation by activating cryptic prophage lytic proteins, causing cell death. (A) Single cell persister resuscitation for wild-type BW25113, the *cas2* mutant, and the Δspacer mutant after 6 hours on 0.4 wt% glucose minimal medium. Black arrows indicate cells that resuscitate, and the scale bar indicates 10 µm. Cells were observed using light microscopy (Zeiss Axio Scope.A1). Representative results from two independent cultures are shown, and tabulated cell numbers are in Table S1. (B) LIVE/DEAD staining of resuscitating persister cells shows the *cas2* and Δspacer mutations cause cell death. DF is dark field, SYTO9 is a membrane permeable stain for nucleic acids (green), and PI is propidium iodide, which is a membrane impermeable stain for the nucleic acids of dead cells (red). Representative results from two independent cultures are shown, and tabulated cell numbers are in Table S2. (C) The *cas2* mutation derepresses cryptic prophage lysis genes *ydfD* (in resuscitating persister cells) and *rzoD* (in both resuscitating persister cells and exponentially-growing cells). Five lytic genes from three cryptic prophages were checked by qRT-PCR: *ydfD* (Qin), *hokD* (Qin), *ypjF* (CP4-57), *essD* (DLP-12), and *rzoD* (DLP-12). Error bars indicate standard deviations from two independent cultures.

### CRISPR-Cas prevents cell death by preventing cell lysis

To explore how deletion of CRISPR-Cas decreases growth (including persister cell resuscitation), we checked for death among resuscitating persister cells of the *cas2* deletion strain using the Live/Dead stain. We found that inactivating CRISPR-Cas leads to a 7-fold increase in death among resuscitating cells (**Fig. 1B, Table S2**). In addition, there were 34-fold more cells termed “ghosts” (Cheng et al., 2014) that lack cytosolic material (**Fig. S5A**), are likely dead, and have intact membranes, so these cells are not stained by the propidium iodide dye. Corroborating these results, there was 11-fold and 5-fold more death for stationary- and exponential-phase cells, respectively, when CRISPR-Cas was inactivated via *cas2* (**Fig. S6, Table S2**). Moreover, for the Δspacer strain, there was 81-fold more cell death in the for stationary-phase cells (**Fig. S6**) and 7-fold more death for resuscitating persister cells (**Fig. 1B, Table S2**). Critically, for the Δspacer strain, waking persister cells clearly shows lysis (**Fig. S5A**), and transmission electron microscopy images confirm this lysis with cytosolic materials seen next to lysed cells (**Fig. S5B**). Unfortunately, this toxicity could not be reversed by CRISPR-Cas induction after placing the complete CRISPR-Cas system on the chromosome under inducible control (BW40114), since this led to more rapid growth when CRISPR-Cas was not induced as a result of the metabolic burden of this complete system; hence, inducing CRISPR-Cas reduced growth (**Fig. S7**). These results indicate inactivating CRISPR-Cas leads to cell death and that the mechanism for cell death is via lysis.

### CRISPR-Cas reduces cryptic prophage lysis gene mRNA

We hypothesized that since CRISPR-Cas systems inhibit some phage before they become lysogens (Edgar and Qimron, 2010; HYPERLINK \l "bookmark11" Goldberg *et al*., 2018), the *E. coli* system may be preventing cell death by repressing certain cryptic prophage genes. To test this hypothesis, we first examined the *E. coli* CRISPR-Cas system for spacers related to the nine cryptic prophages. *E. coli* K-12 CRISPR array located next to the *cas* operon contains 13 spacers (Militello and Lazatin, 2017; HYPERLINK \l "bookmark30" Pougach *et al*., 2010), each containing 32 or 33 nt (Brouns et al., 2008). Between the 14 29-nt repeat sequences (5’-GTGTTCCCCGCATCAGCGGGGAfTAAACCG), we found that four of the 13 spacers contain 7 to 16 nt of perfect matches to seven of the nine cryptic prophages (DLP-12, CPS-53, CP4-6, Rac, Qin, CP4-57, and e14) (**Fig. 2A**) based on basepairing matches with cryptic prophage mRNA (**Fig. S8**), indicating putative binding to cryptic prophage mRNA. Critically, spacer 12 encodes crRNA with the potential to bind mRNA from DLP-12 (*rzoD, ybcN*), Rac (*stfR*), Qin (*stfQ*), and CP4-57 (*alpA*) (**Fig. S8**). In general, spacer lengths vary from 21 to 72 nt (Barrangou and Marraffini, 2014) with perfect complementarity of 6 to 12 nt (Shmakov *et al*., 2017). Together, these results suggest CRISPR-Cas potentially regulates the *E. coli* cryptic prophages. Note that the second CRISPR array in *E. coli* K-12 in between genes *ygcE* and *queE* lacks spacers with sequences that matched the cryptic prophages.

**Figure 2.**
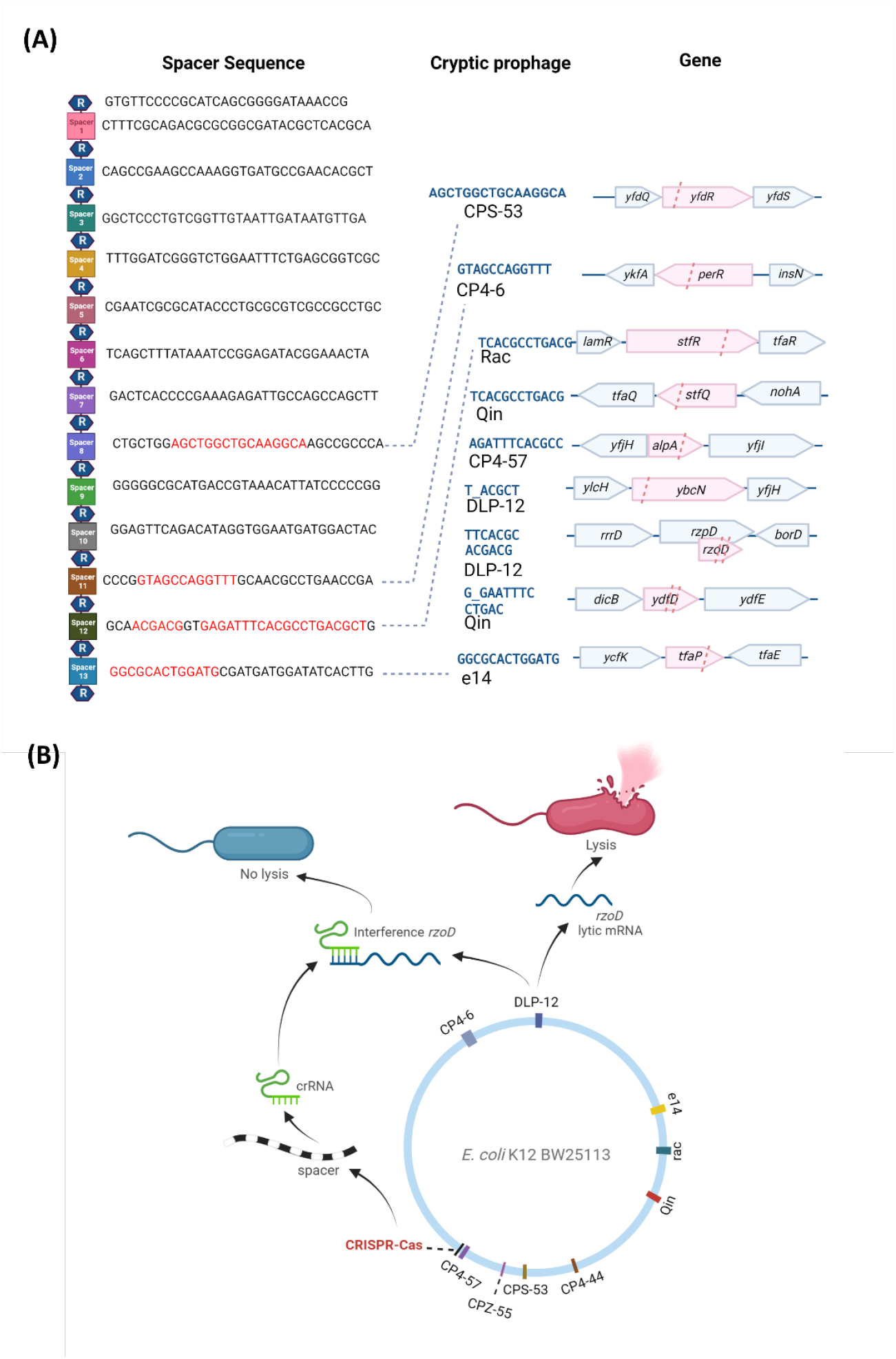
*E. coli* CRISPR-Cas spacer sequences and lytic gene inhibition mechanism. **(A)** The 14 repeat (R, hexagon) and 13 spacer (squares) sequences of the CRISPR-Cas system (from the *iap* to *cas2* part of the *E. coli* genome) showing the cryptic prophage spacer matches (red text) and prophage DNA protospacer sequences (blue text) which includes matches to seven of the nine cryptic prophages (CPS-53, CP4-6, Rac, Qin, CP4-57, DLP-12, and e14). Matches indicate mRNA binding to spacer sequences (**Fig. S8**). Pink highlight and pink dashed lines indicate the spacer positions relative to the cryptic prophage genes. **(B)** Schematic for a hypothetical mechanism by which CRISPR-Cas controls cryptic phage lytic genes.

The presence of the cryptic prophage-related targeting spacers suggests that *E. coli* CRISPR-Cas may be preventing expression of cryptic prophage genes and, given the cell phenotype, is suggestive of prophage-encoded lethal genes; hence, we checked the mRNA levels of the cryptic prophage lysis genes encoded by these seven cryptic prophages with spacer matches. Specifically, transcription of *ydfD* (Qin), *hokD* (Qin), *ypjF* (CP4-57), *essD* (DLP-12), and *rzoD* (DLP-12) was checked via quantitative reverse transcription polymerase chain reaction (qRT-PCR). We found that inactivation of CRISPR-Cas via the *cas2* deletion results in a 35-fold increase in *ydfD* mRNA and a 39-fold increase in *rzoD* mRNA in resuscitating persister cells (**Fig. 1C**). RzoD is a putative DLP-12 lysis lipoprotein that we previously showed was toxic through its interaction with toxin Hha (García Contreras et al., 2008) of the Hha/TomB TA system (Marimon et al., 2016). YdfD of Qin cryptic prophage has been shown to lyse cells when induced (Masuda et al., 2016). To confirm that YdfD and RzoD are toxic, we induced production of both proteins and found they are indeed toxic (**Fig. S9**). Moreover, deleting *cas2* to inactivate CRISPR-Cas in the same host that lacks all nine cryptic prophages (delta9) (Wang *et al*., 2010) has no effect on growth (**Fig. S10**) and does not cause lysis (**Fig. S6**). Hence, these results show CRISPR-Cas represses at least two *E. coli* cryptic prophage proteins RzoD and YdfD that can reduce cell growth.

### Excision of DLP-12 is not regulated by CRISPR-Cas

Since inactivation of CRISPR-Cas leads to derepression of the DLP-12 *rzoD* lysis gene, we checked for increased DLP-12 excision with the *cas2* deletion strain using qPCR. We found the *cas2* deletion has little impact on DLP-12 excision (**Table S3**). Since the DLP-12 toxin gene *essD* is not derepressed, the effect is not due to excision of the prophage. We also tested the effect of CRISPR-Cas on two other cryptic prophages with significant excision (CP4-57 and e14) (Wang *et al*., 2010), and also found no effect of deleting *cas2* (**Table S3**). Hence, CRISPR-Cas does not affect cryptic prophage excision.

### CRISPR-Cas regulates lytic gene mRNA levels

Since CRISPR-Cas does not affect cryptic prophage excision and is unlikely to degrade cryptic prophage DNA as this would be lethal (Edgar and Qimron, 2010; HYPERLINK \l "bookmark11" Goldberg *et al*., 2018), we reasoned that CRISPR-Cas, via crRNA, must be interfering with expression of the mRNA of the lytic genes to prevent production of the lytic proteins. To test this hypothesis, we investigated whether *rzoD* and *ydfD* transcript levels are increased in the *cas2* and Δspacer strain relative to the wild-type after inhibiting transcription via rifampicin. If CRISPR-Cas prevents lytic gene expression by inhibiting lytic transcripts via crRNA, then inactivating CRISPR-Cas should lead to increased *rzoD* and *ydfD* mRNA. In accordance with this hypothesis, *rzoD* mRNA was increased 320% in the Δspacer mutant and 50% in the *cas2* mutant. Moreover, as expected, inactivating CasE increased cell lysis 49-fold compared to the wild-type strain (**Fig. S6, Table S2)**, likely by preventing CasE from properly forming crRNA from pre-crRNA; hence, there is less crRNA to inhibit the cryptic prophage mRNA from toxic genes. Similarly, preventing spacer RNA formation by inactivating Cas2 led to a 10-fold increase in CRISPR array RNA levels in resuscitating persister cells as assayed by qRT-PCR, likely due to a lack of processing of the CRISPR array into crRNA (**Table S3B**). Finally, deleting only spacer #12 (which encodes crRNA targeting three separate regions in the cryptic prophage mRNA of CP4-57, Rac, and Qin, **Fig. S8**), in a markerless fashion, inhibits cell growth in the stationary phase (**Fig. S10**) due to cell lysis (**Fig. S6, Table S2**) to roughly the same extent as deleting all 13 spacers. In contrast, deleting spacer #3 in a markerless fashion, which does not encode crRNA with matches to cryptic prophage mRNA, has little effect on overall growth (**Fig. S10**), but did increase cell death (**Fig. S6, Table S2**), which indicates it does serve some role. Therefore, CRISPR-Cas prevents cryptic prophage lytic gene expression likely by interfering with the transcripts from these genes (**Fig. 2B**), specifically through spacer #12.

## 3. DISCUSSION

Our results reveal a new (non-canonical) role for CRISPR-Cas systems: regulation of phage fossils. The evidence for this includes that inactivating CRISPR-Cas by deleting *cas2* (i) reduces growth by 40%, (ii) nearly eliminates resuscitation from the persister state (**Fig. 1**), (iii) causes ghost cell formation and cell death (**Fig. S5**), (iv) increases mRNA levels of the cryptic prophage lysis genes *ydfD* and *rzoD*, and (v) increases CRISPR array levels by preventing processing into crRNA. Corroborating these results, the *cas2* deletion has no effect in a strain which lacks cryptic prophage genes and eliminating the CRISPR array with prophage-matching spacers in the wild-type strain also reduced the growth rate, eliminated persister cell resuscitation, resulted in cell death via lysis, and activated *rzoD*. Furthermore, deleting only prophage-matching spacer #12 similarly inhibited growth and caused cell lysis. Given that Cas2 is not required for CRISPR interference, it is likely that the phenotypes produced upon deleting *cas2* are the result of a polar mutation (**Fig. S3**) that inactivates CRISPR-Cas by altering production/processing of the spacers.

Since there is no change in excision of DLP-12 upon inactivating CRISPR-Cas, our results suggest the mechanism for regulating the lysis genes *ydfD* and *rzoD* of the cryptic prophages is via CRISPR-Cas RNA binding to the cryptic prophage mRNA (analogous to RNA interference as seen previously in procaryotic systems with CRISPR-Cas (Mohanraju et al., 2022)) rather than cleaving excised DNA or directly cleaving the mRNA. Our proposed RNAi mechanism has precedent since a previous bioinformatics analysis of 230 phages suggested CRISPR-Cas functions in *E. coli* via endogenous gene expression rather than acting as an immune system (Bozic et al., 2019). In support of this, the excision rates of the cryptic prophages are so low that there may be insufficient excised DNA to cleave (1 excision in 10^6^ cells for CP4-57) (Wang *et al*., 2010), and a class I-F CRISPR-Cas system in *P. aeruginosa* uses crRNA to regulate endogenous LasR mRNA to reduce the pro-inflammatory response (Li et al., 2016). Moreover, *E. coli* crRNA have been shown to be activated during infection in mice (Li *et al*., 2016), and the *Salmonella* spp. system is identical to the *E. coli* one (e.g., the repeats are the same), but the spacers are different, which suggests it, too, is for endogenous gene expression. Critically, it has been demonstrated *in vitro* that that the *E. coli* Cascade binds ssRNA complementary to crRNA (Jore et al., 2011). Therefore, there is ample precedence for our proposed RNA interference mechanism.

Although we identified spacer #12 sequences that probably bind and interfere with the cryptic prophages Qin 12 (including *ydfD*) and DLP-12 (including *rzoD*) mRNA, there may be another level of regulation that remains to be identified. For example, we show a spacer #12 region that would bind *alpA* mRNA in cryptic prophage CP4-57, and AlpA is a regulator that impacts persister resuscitation by sensing phosphate nutrients (Song et al., 2021).

This new function for CRISPR-Cas acting on cryptic prophages is likely general and may explain why many species appear to have inactive CRISPR-Cas systems as was previously thought for *E. coli* (Shmakov *et al*., 2017); i.e., instead of protecting cells from external phages, CRISPR-Cas systems may also control resident cryptic prophages which are prevalent. As additional evidence, we found matches in CRISPR-Cas spacers not only for *E. coli* K-12 (**Fig. 2A**) but also for *E. coli* O157:H7, *Salmonella* spp., and *K. pneumoniae* (**Fig. S11**). Critically, our results provide the first example where it is beneficial for the host to have an active CRISPR-Cas system that targets *inactive* integrated phages (i.e., cryptic prophages) since previous reports show targeting *active* temperate phages is deleterious; i.e., either the cell dies or the phage is lost (Edgar and Qimron, 2010; HYPERLINK \l "bookmark11" Goldberg *et* al., 2018; Rollie et al., 2020). Since *E. coli* cryptic prophages like *rac* have been present in its genome for 4.5 million years (Perna et al., 2001), the active *E. coli* K-12 CRISPR-Cas system is stable with the cryptic prophages; in fact, there has been little change in the *E. coli* spacers for at least 42,000 years (Savitskaya et al., 2017).

Our results also indicate that, although the cryptic prophages are stable and the cell makes use of the genetic tools encoded by its former foe to combat myriad stresses (Wang *et al*., 2010) and to sense nutrients prior to exiting the persister state (Song *et al*., 2021), the source of these tools must be elegantly regulated by CRISPR-Cas since they often harbor deleterious membrane lysis proteins like YdfD and RzoD. Similarly, host Rho has been found recently to silence cryptic prophage toxin/antitoxin systems through transcription termination (Hafeezunnisa et al., 2021), and H-NS silences cryptic prophages through 65 binding sites (Ishihama and Shimada, 2021). Therefore, phages may be captured by the host, but they must be tamed, and this now includes repression by CRISPR-Cas. These results also suggest a role for CRISPR-Cas in gene regulation beyond self-defense.

## 4. MATERIALS AND METHODS

### Bacteria and growth conditions

Bacteria (**Table S4**) were cultured routinely in lysogeny broth (Bertani, 1951) at 37°C, while M9 glucose (0.4 wt%) (Rodriguez and Tait, 1983) was used to assay the growth rate and to resuscitate persister cells. pCA24N-based plasmids (Kitagawa et al., 2005) were retained in overnight cultures via chloramphenicol (30 µg/mL), and kanamycin (50 µg/mL) was used for deletion mutants, where applicable. The BW25113 *cas2* and spacer deletion were confirmed by PCR (primers shown in **Table S5**). Spacers 3 and 12 were deleted from BW25113, and *cas2* was deleted from the strain that lacks cryptic prophages (Δ9) using the method of Datsenko and Wanner (Datsenko and Wanner, 2000) using plasmids pKD4 with the primers in **Table S5**; kanamycin resistance was removed using pCP20 (Datsenko and Wanner, 2000).

### Persister cells

Exponentially-growing cells (turbidity of 0.8 at 600 nm) were converted nearly completely to persister cells (Kim *et al*., 2018; HYPERLINK \l "bookmark18" Kwan *et al*., 2013) by adding rifampicin (100 µg/mL) for 30 min to stop transcription, centrifuging, and adding LB with ampicillin (100 µg/mL) for 3 h to lyse non-persister cells. To remove ampicillin, cells were washed twice with 0.85% NaCl then re-suspended in 0.85% NaCl. Persister concentrations were enumerated via a drop assay (Donegan et al., 1991).

### Single-cell persister resuscitation

Persister cells (5 µL) were added to 1.5% agarose gel pads containing M9 glucose (0.4 wt%) medium (Rodriguez and Tait, 1983), and singlecell resuscitation was visualized at 37°C via a light microscope (Zeiss Axio Scope.A1, bl_ph channel at 1000 ms exposure). For each condition, at least two independent cultures were used with 150 to 300 individual cells assessed per culture.

### Membrane integrity assay

To determine membrane integrity, the persister cells were analyzed with the LIVE/DEAD BacLight Bacterial Viability Kit (Molecular Probes, Inc., Eugene, OR, catalog number L7012). The fluorescence signal was analyzed via a Zeiss Axioscope.A1 using excitation at 485 nm and emission at 530 nm for green fluorescence and using excitation at 485 nm and emission at 630 nm for red fluorescence.

### qRT-PCR

To quantify transcription from the cryptic prophage lytic genes, RNA was isolated from persister cells that were resuscitated by adding M9 glucose (0.4 wt%) medium for 10 min and from exponential cells grown to a turbidity of 0.8. For quantifying transcription from the CRISPR array, RNA was isolated from persister cells that were resuscitated by adding M9 glucose (0.4 wt%) medium for 10 min. Samples were cooled rapidly using ethanol/dry ice in the presence of RNA Later. RNA was isolated using the High Pure RNA Isolation Kit (Roche). The following qRT-PCR thermocycling protocol was used with the iTaq™ universal SYBR^®^ Green One-Step kit (Bio-Rad): 95 ºC for 5 min; 40 cycles of 95 ºC for 15 s, 60 ºC for 1 min for two replicate reactions for each sample/primer pair. The annealing temperature was 60ºC for all primers (**Table S5**).

### qPCR

To quantify prophage excision and the levels of DNA flanking the CRISPR-Cas cleavage sites, total DNA (100 ng) was isolated from exponentially-growing and persister resuscitating cells using an UltraClean Microbial DNA Isolation Kit (Mo Bio Laboratories). Excised cryptic prophage was quantified using primers for each prophage excisionase (**Table S5**) that only yield a PCR product upon prophage excision, and the relative amount of each target gene was determined using reference gene *purM*. The level of cryptic prophage flanking the CRISPR-Cas cleave site was quantified using primers that flank each site (**Table S5**). The qPCR reaction performed using CFX96 Real Time System. The reaction and analysis was conducted using the StepOne Real-Time PCR System (Bio-Rad).

### Transmission electron microscopy

For transmission electron microscopy (TEM), the samples were prepared from persister cells that were resuscitated by adding M9 glucose (0.4%) medium for 10 min then washed with 0.85% NaCl. The samples were fixed with buffer (2.5% glutaraldehyde in 0.1M cacodylate buffer, pH 7.4). The negative staining was performed with 2% uranyl acetate for 1 hr, then dehydrated. The sectioned specimens were stained again with uranyl acetate and lead citrate after dehydration and embedded with resin. The TEM image was observed using a Hitachi (H-7605) instrument.

## ACKNOWLEDGEMENTS

This work was supported by funds derived from the Biotechnology Endowed Professorship at the Pennsylvania State University for TKW and from the National Research Foundation of Korea (NRF) grant from the Korean Government (NRF-2020R1F1A1072397) for SYS. We appreciate the feedback of Joy Muthami. The authors have no conflicts of interest.

## Supporting Information

**Table S1.** Inactivating CRISPR-Cas eliminates persister cell resuscitation on glucose agarose gel pads. Single persister cells were observed using light microscopy (Zeiss Axio Scope.A1). The total number and waking number of persister cells are shown after 6 hours on 0.4 wt% glucose minimal medium. Fold-change in waking is relative to BW25113. These results are the combined observations from two independent experiments (independent culture results separated by “/”), and standard deviations are shown. Sample microscope images are shown in **Fig. 1**.

|  | **Total**<br>**cells** | **Waking**<br>**cells** | **% waking** | **Fold-**<br>**change** |
| --- | --- | --- | --- | --- |
| BW25113 | 504/382 | 199/248 | 52 ± 18 | 1 |
| *Δcas2* | 516/677 | 6/15 | 1.7 ± 0.7 | -30.9 |
| *Δspacer* | 323/174 | 10/7 | 3.6 ± 0.7 | -14.6 |

**Table S2.** Inactivating CRISPR-Cas causes cell death and ghost cell formation. For resuscitated cells, persister cells were washed with PBS twice, resuscitated by M9 0.4% glucose for 10 min and stained with LIVE/DEAD reagents. Exponential cells were grown to a turbidity of 0.8 (at 600 nm) and stationary cells were grown to turbidity of 2.0. The ghost cells in the persister population were visualized using a Zeiss Axioscope.A1microscope. The results are the combined observations from two independent experiments (independent culture results separated by “/”). The microscope images are shown in **Fig. 1AB** and **Fig. S6**.

|  | Strains | Total cells | Dead cells | Ghost cells | % Ghost cells | fold-change ghost cells | % dead | fold-change dead cells |
| --- | --- | --- | --- | --- | --- | --- | --- | --- |
| **Resuscitated** | **BW25113** | 52/115 | 2/7 | 0/1 | 0.6 | 1 | 5.4 | 1 |
| | $\Delta cas2$ | 142/106 | 54/38 | 38/13 | 20.6 | 34 | 37.1 | 6.9 |
| | $\Delta spacer$ | 161/221 | 76/76 | - | - | - | 37.8 | 7.0 |
| **Exponential** | **BW25113** | 153/147 | 0/2 | - | - | - | 0.7 | 1 |
| | $\Delta cas2$ | 59/40 | 1/2 | - | - | - | 3.4 | 4.9 |
| | $\Delta spacer$ | 95/146 | 1/5 | - | - | - | 2.1 | 3.15 |
| **Stationary** | **BW25113** | 492/123 | 1/0 | - | - | - | 0.1 | 1 |
| | $\Delta cas2$ | 1136/173<br>6 | 14/18 | - | - | - | 1.13 | 11.2 |
| | $\Delta spacer$ | 200/113 | 19/8 | - | - | - | 8.26 | 81.3 |
| | $\Delta casE$ | 208/175 | 9/10 | - | - | - | 5.02 | 49.4 |
| | $\Delta 9 \Delta cas2$ | 833/1541 | 10/13 | - | - | - | 1.02 | |
| | $\Delta spacer3$ | 136/297 | 8/73 | - | - | - | 15.23 | |
| | $\Delta spacer12$ | 513/396 | 213/92 | - | - | - | 32.38 | |

**Table S3.** CRISPR-Cas does not affect cryptic prophage excision in stationary cells but produces the CRISPR array. (**A**) Fold changes in excision are relative to the wild-type strain as determined by qPCR. (**B**) Fold changes in the CRISPR array are relative to the wild-type strain as determined by qRT-PCR (the Δspacer strain was used as a negative control). Cycle numbers (C_t_) are indicated for each sample including that for the target genes as well as that of the house-keeping gene, *purM*, which was used to normalize the data. *cas2* served to inactivate crRNA production due to a polar mutation. Fold changes in the transcription of various targets (i.e., cryptic prophage CP4-57, e14, DLP-12, Spacer3, and Spacer12) with and without *cas2*^+^ (*cas2*^+^/*cas2*^-^) were calculated as described earlier (Pfaffl, 2001):

**(A) qPCR**
| Gene | *purM* |  | CP4-57 |  | e14 |  | DLP-12 |  |
| --- | --- | --- | --- | --- | --- | --- | --- | --- |
| Strain | WT | *cas2* | WT | *cas2* | WT | *cas2* | WT | *cas2* |
| $C_t$ | 9.30<br>± 0.13 | 9.70<br>± 0.18 | 30.45<br>± 1.73 | 30.37<br>± 0.68 | 16.30<br>± 0.06 | 17.45<br>± 0.51 | 30.58<br>± 0.49 | 30.05<br>± 0.5 |
| $\Delta C_t$ | | | 21.16<br>± 1.73 | 20.67<br>± 0.70 | 7.00<br>± 0.15 | 8.16<br>± 0.54 | 21.29<br>± 0.51 | 20.35<br>± 0.53 |
| $\Delta\Delta C_t$ | | | | -0.48<br>± 0.7 | | 1.16<br>± 0.54 | | -0.94<br>± 0.53 |
| fold |  |  |  | 1.40 |  | -2.23 |  | 1.92 |

**(B) qRT-PCR**
| Gene | *purM* |  |  | Spacer3 |  |  | Spacer12 |  |  |
| --- | --- | --- | --- | --- | --- | --- | --- | --- | --- |
| Strain | WT | *cas2* | spacer | WT | *cas2* | spacer | WT | *cas2* | spacer |
| $C_t$ | 25.3<br>± 0.5 | 22.2<br>± 0.5 | 22.0<br>± 0.2 | 30.68<br>± 1.73 | 24.35<br>± 0.93 | - | 33.60<br>± 1.20 | 27.12<br>± 1.22 | - |
| $\Delta C_t$ | | | | 5.37<br>± 1.08 | 2.19<br>± 1.1 | - | 8.28<br>± 1.31 | 4.96<br>± 1.32 | - |
| $\Delta\Delta C_t$ | | | | | -3.20<br>± 1.51 | - | | -3.33<br>± 1.86 | - |
| fold |  |  |  |  | 9.11 | -∞ |  | 10.02 | -∞ |

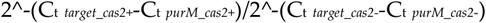

**Table S4.** *E. coli* bacterial strains and plasmids utilized.

| Strains and Plasmids | Features | Source |
| --- | --- | --- |
| **Strains** |  |  |
| BW25113 | *rrnB3 ΔlacZ4787 hsdR514 Δ(araBAD)567 Δ(rhaBAD)568 rph-1* | (Baba et al., 2006) |
| BW25113 *Δcas1* | *Δcas1, ΩKm<sup>R</sup>* | (Baba et al., 2006) |
| BW25113 *Δcas2* | *Δcas2, ΩKm<sup>R</sup>* | (Baba et al., 2006) |
| BW25113 *Δcas2 ΔKm<sup>R</sup>* | *Δcas2, ΔKm<sup>R</sup>* | this work |
| BW25113 *Δcas3* | *Δcas3, ΩKm<sup>R</sup>* | (Baba et al., 2006) |
| BW25113 *ΔcasA* | *ΔcasA, ΩKm<sup>R</sup>* | (Baba et al., 2006) |
| BW25113 *ΔcasB* | *ΔcasB, ΩKm<sup>R</sup>* | (Baba et al., 2006) |
| BW25113 *ΔcasC* | *ΔcasC, ΩMr* | (Baba et al., 2006) |
| BW25113 *ΔcasD* | *ΔcasD, ΩKm<sup>R</sup>* | (Baba et al., 2006) |
| BW25113 *ΔcasE* | *ΔcasE, ΩKm<sup>R</sup>* | (Baba et al., 2006) |
| BW25113 *Δspacer*<br>(also called BW39292) | *Δ(13 spacer region), ΩKm<sup>R</sup>* | (Pougach et al., 2010) |
| BW40114 | *placuv5::cas3 paraB8p::casABCDE12spacers* | (Datsenko et al., 2012) |
| BW25113 *Δspacer3* | *Δ(spacer region 3), ΔKm<sup>R</sup>* | this work |
| BW25113 *Δspacer12* | *Δ(spacer region 12), ΔKm<sup>R</sup>* | this work |
| BW40114 | BW25113 *lacUV5::cas3 araB8p::casABCDE12* | (Datsenko et al., 2012) |
| BW15113 *Δ9* | BW15113 lacking all 9 cryptic prophages | (Wang et al., 2010) |
| BW15113 *Δ9 cas2* | BW15113 *Δ9 cas2, ΔKm<sup>R</sup>* | this work |
| *ssrA* | *ssrA, ΩKm<sup>R</sup>* | (Wang et al., 2009) |
| **Plasmids** |  |  |
| pCA24N | *Cm<sup>R</sup>; lacI<sup>q</sup>* | (Kitagawa et al., 2005) |
| pCA24N_*cas2* | *Cm<sup>R</sup>; lacI<sup>q</sup>, P<sub>T5-lac</sub>::cas2<sup>+</sup>* | (Kitagawa et al., 2005) |
| pKD4 | *FRT::Kan<sup>R</sup>::FRT, Amp<sup>R</sup>* | (Datsenko and Wanner, 2000) |
| pCP20 | *FLP synthase, Amp<sup>R</sup>* | (Datsenko and Wanner, 2000) |

**Table S5.**
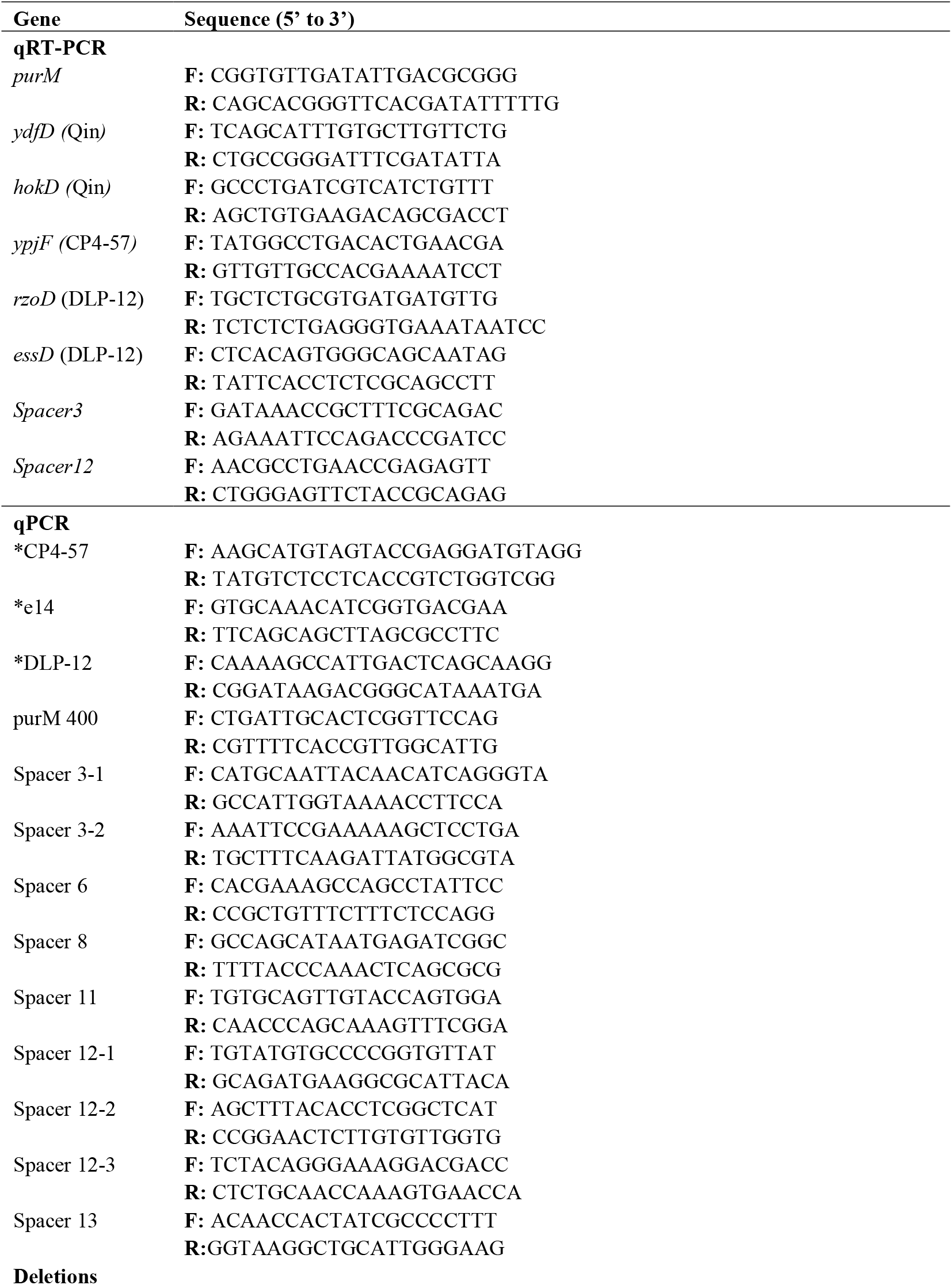
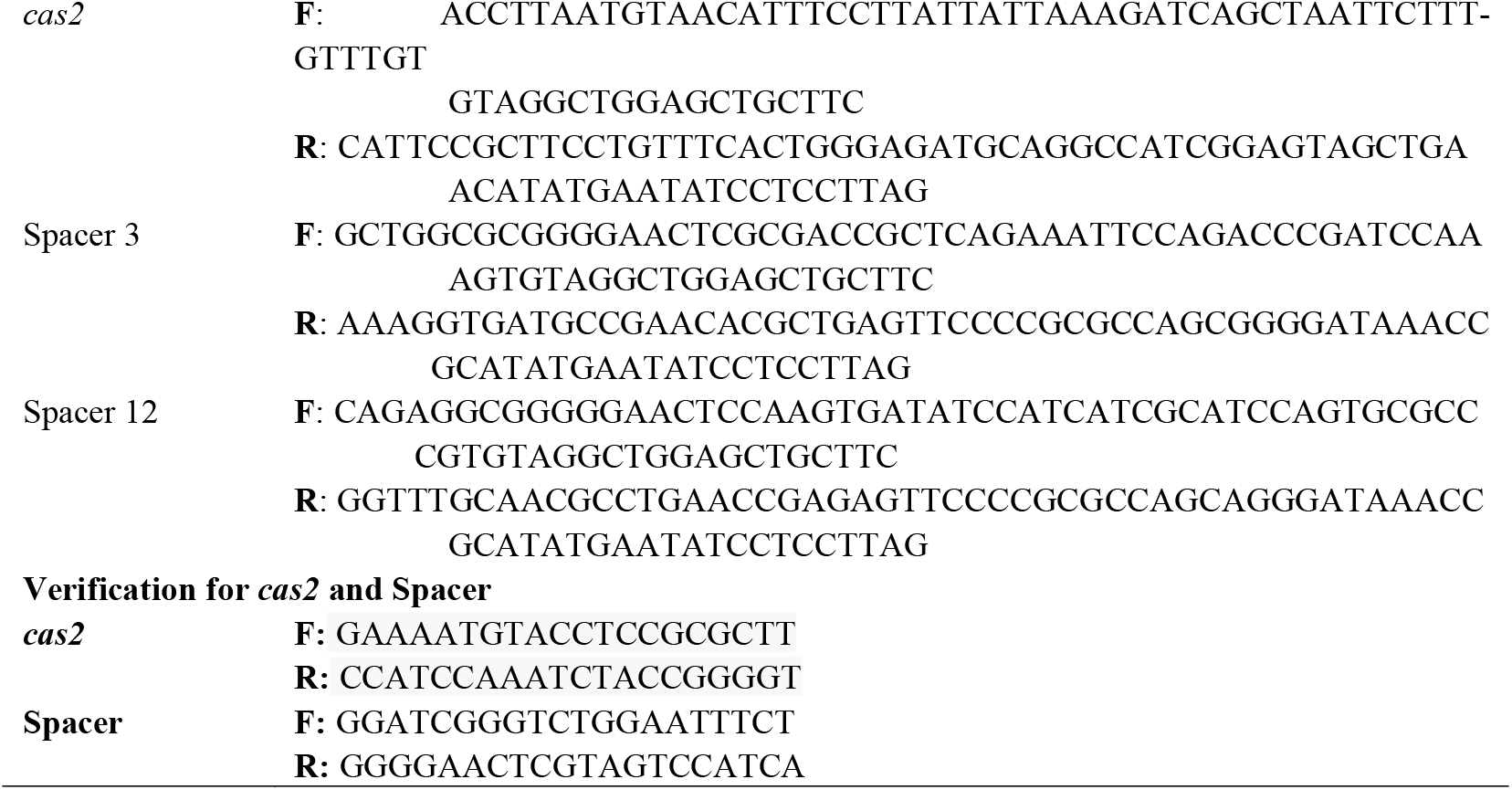
Primers used in this study for qRT-PCR and qPCR. * indicates excision primers.

**Supplementary Figure 1.**
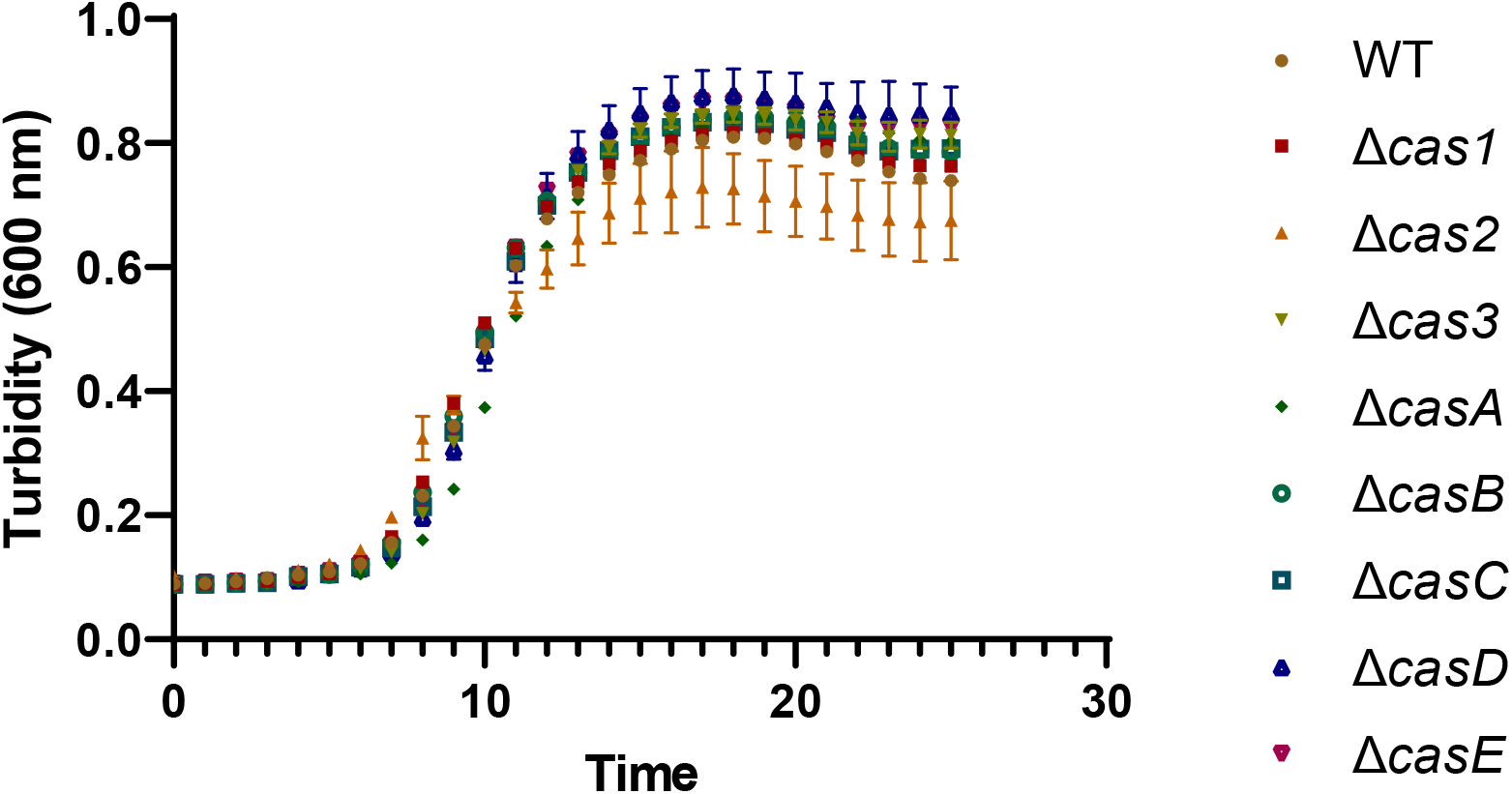
Growth curves for Δ*cas1*, Δ*cas2, cas3*, Δ*casA*, Δ*casB*, Δ*casC*, Δ*casD*, and Δ*casE* in M9 glucose medium at 37°C. Turbidity at 600 nm shown, and error bars indicate standard deviations from two independent cultures.

**Supplementary Figure 2.**
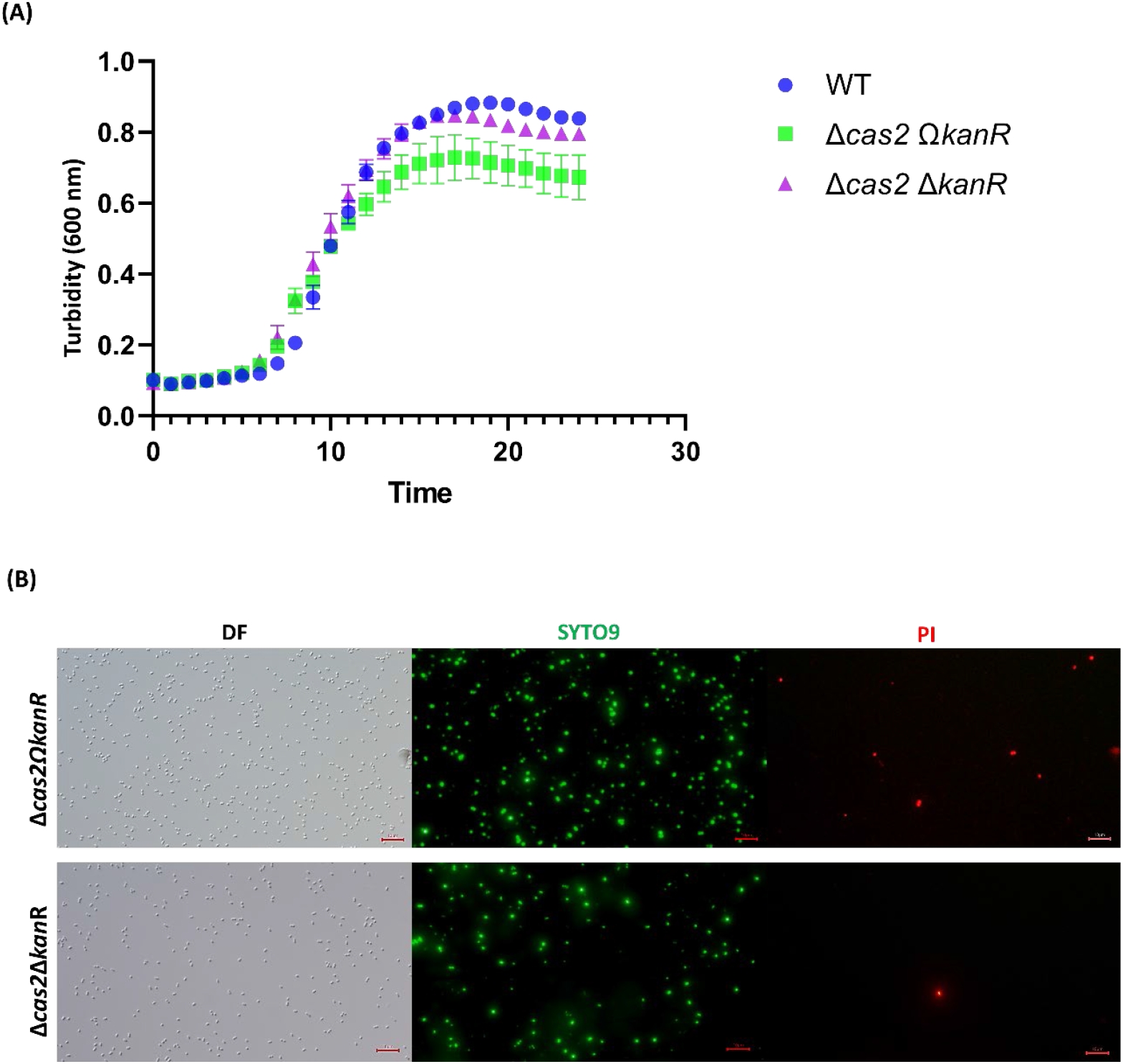
Removing the kanamycin marker from the *E. coli* Δ*cas2* Ω*kan*^*R*^ strain to create the Δ*cas2* Δ*kan*^*R*^ strain **(A)** increased growth in M9 glucose medium at 37°C (turbidity at 600 nm shown and error bars indicate standard deviations from two independent cultures) and **(B)** reduced toxicity via the LIVE/DEAD assay for stationary cells showing the polar mutation due to the kanamycin marker in the Δ*cas2* Ω*kan*^*R*^ strain inhibits spacer formation, which results in reduced growth and increased toxicity. Representative images shown from two independent cultures.

**Supplementary Figure 3.**
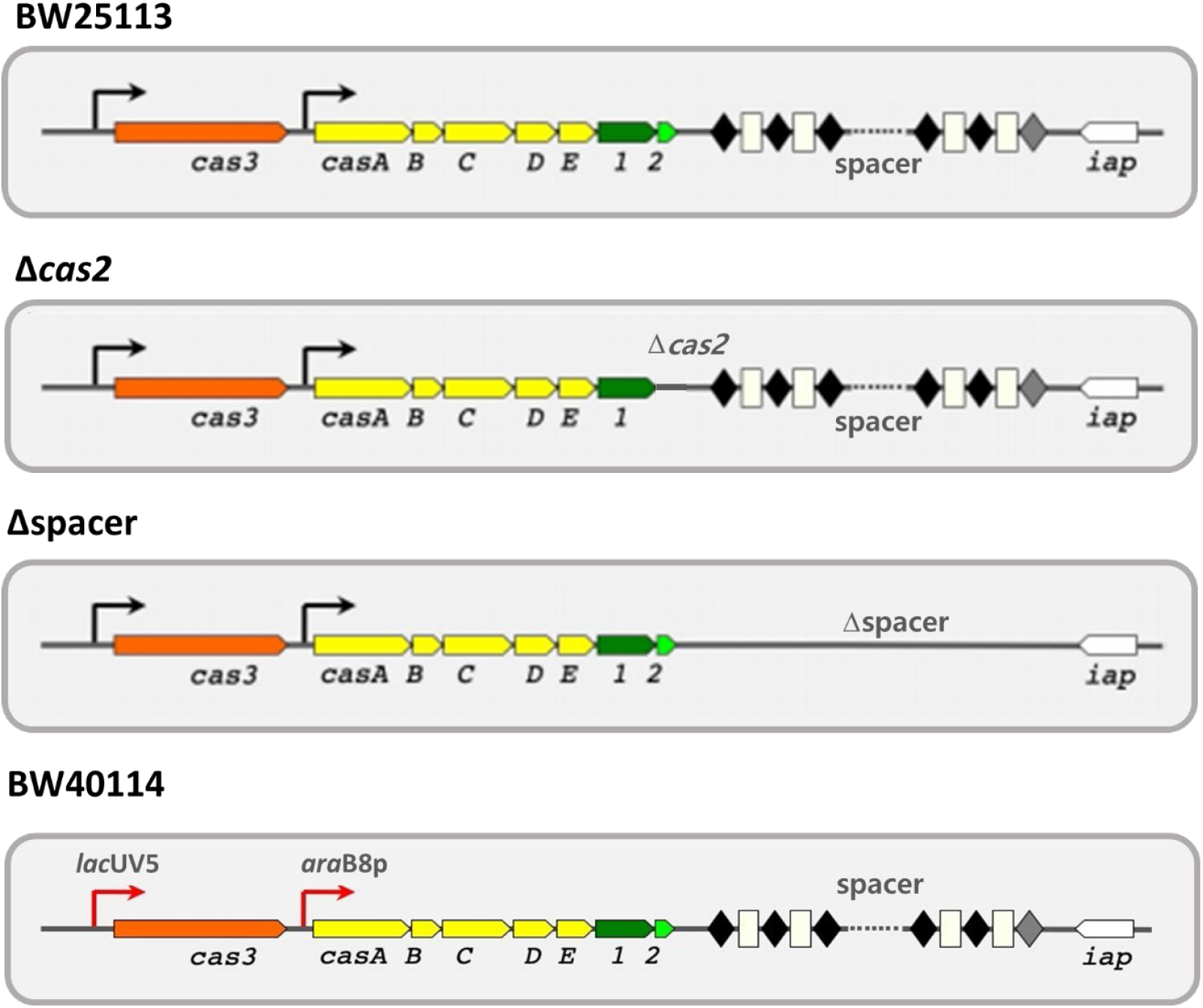
Schematics of the CRISPR-Cas strains used in this study. All the strains are derivatives of isogenic host, BW25113 (see **Table S4**).

**Supplementary Figure 4.**
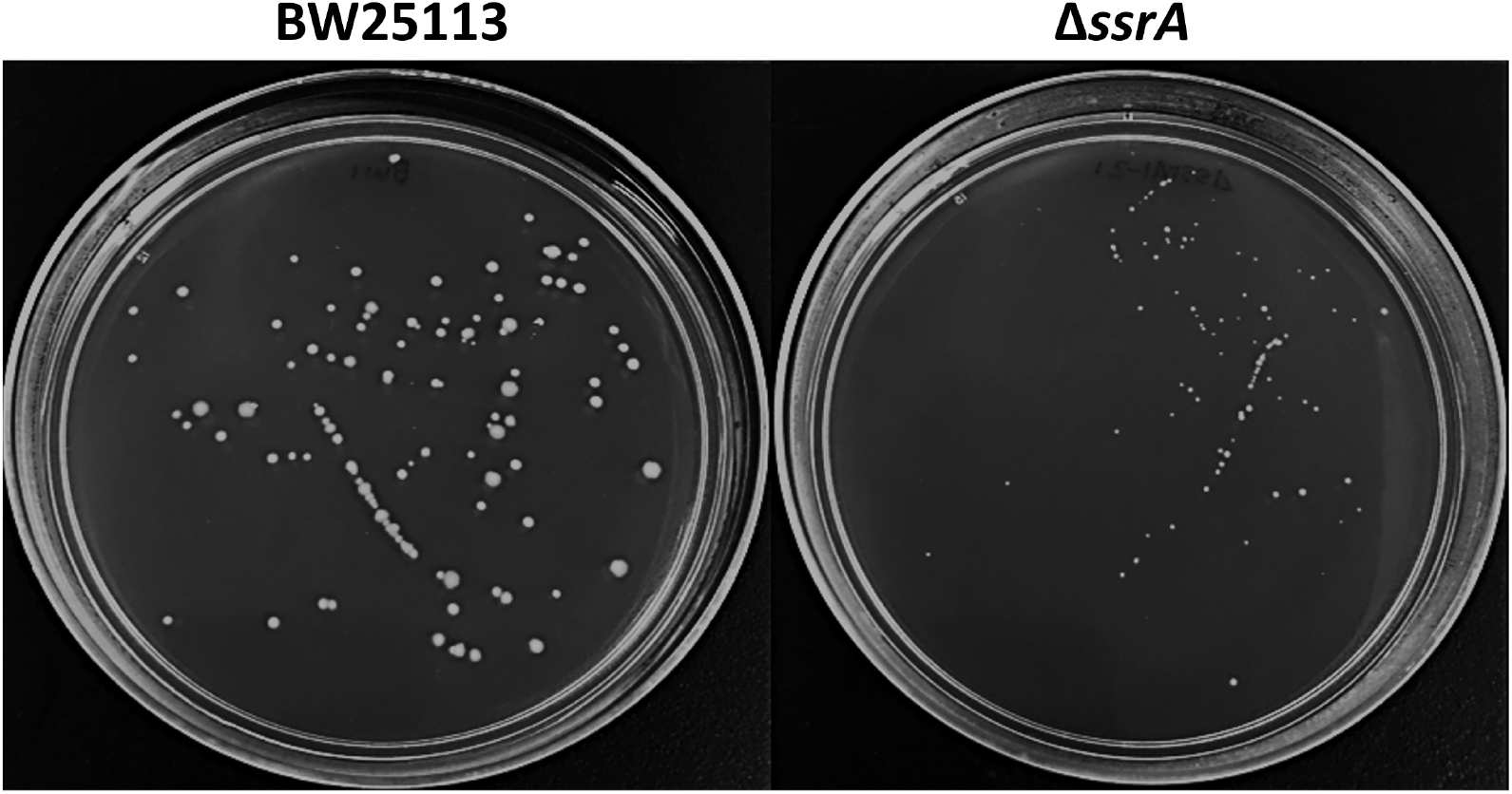
As a control for growth rate and resuscitation, the *ssrA* mutation was checked and found to have no effect on persister resuscitation on LB plates although it grows 22% more slowly than the wild-type strain (1.18 ± 0.07/h vs. 1.49 ± 0.03/h, respectively). The number of *ssrA* persisters that resuscitated is 101 ± 2 vs. 100.0 ± 0.1 for the wild-type. Plates were incubated for 24 h at 37°C, and representative images shown from two independent cultures.

**Supplementary Figure 5.**
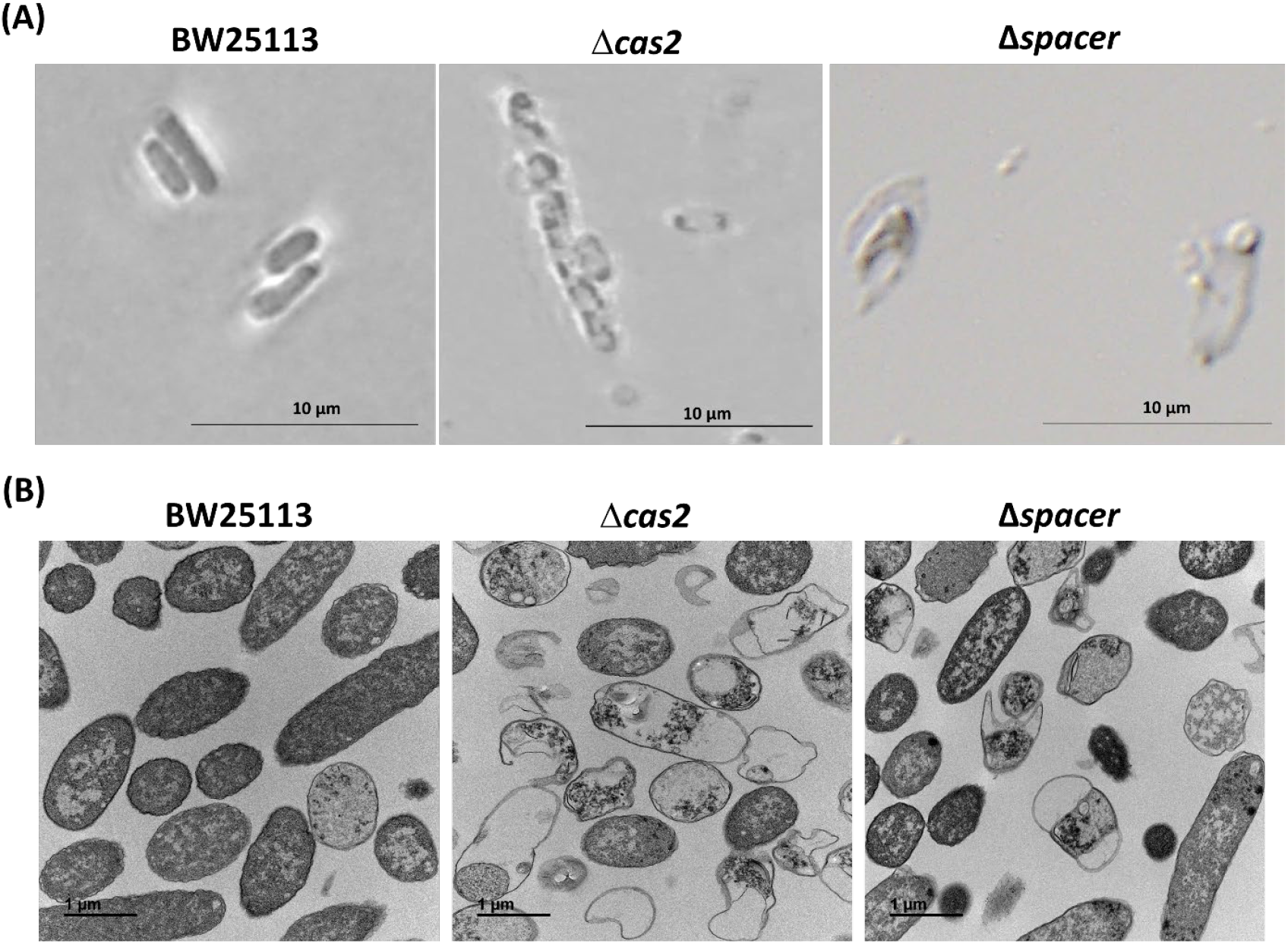
CRISPR-Cas prevents cell lysis induced by cryptic phages. (**A**) Single cell persister waking of BW25113 Δ*cas2* and the Δspacer mutant on M9 0.4% glucose agar plates incubated at 37°C for 4 hours. The scale bar indicates 10µm. (**B**) TEM image for persister waking of BW25113 Δ*cas2* and the Δspacer mutant. Persister cells were resuscitated by M9 0.4% glucose for 10 min. One representative image from two independent cultures is shown. The scale bar indicates 1 µm.

**Supplementary Figure 6.**
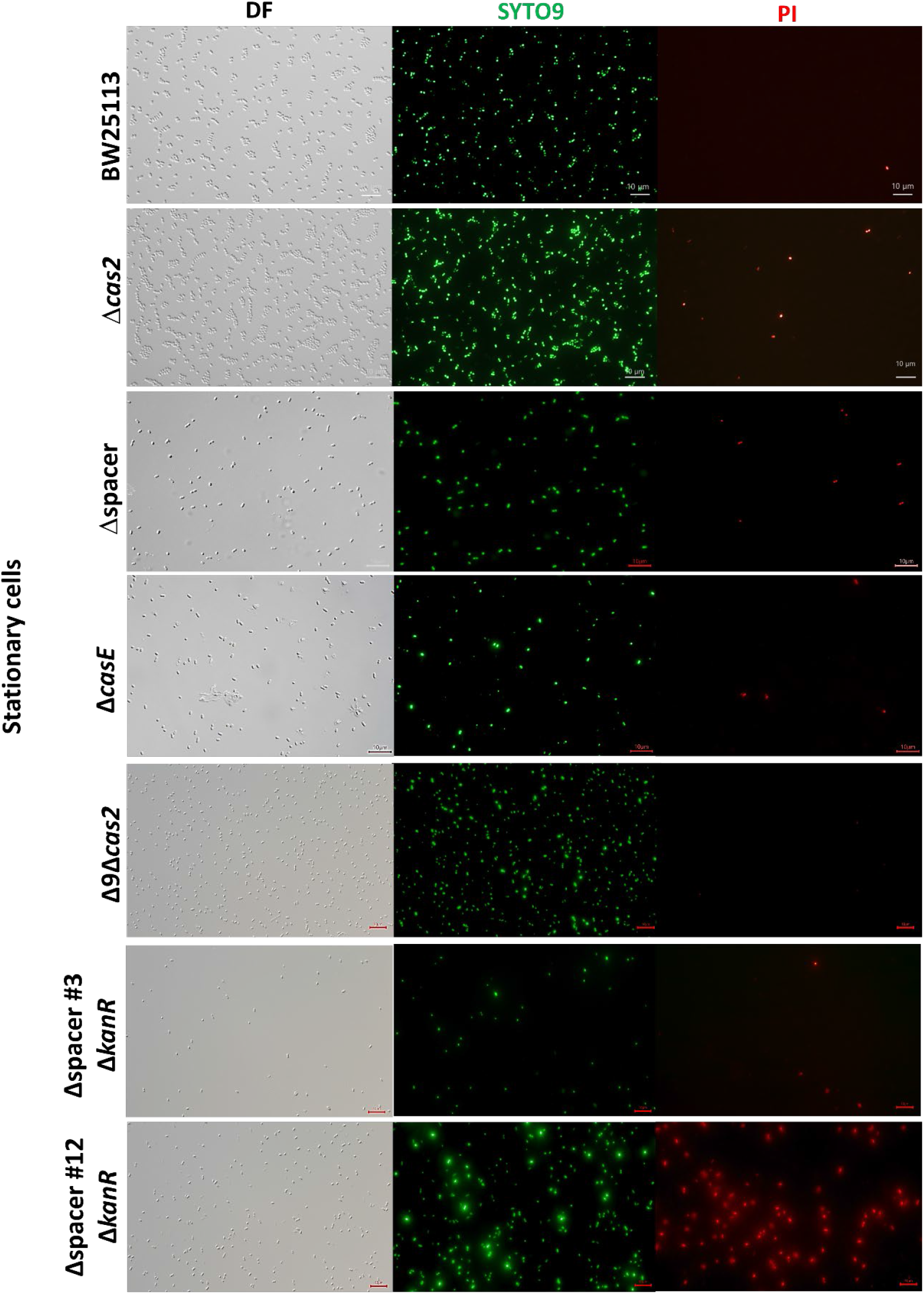
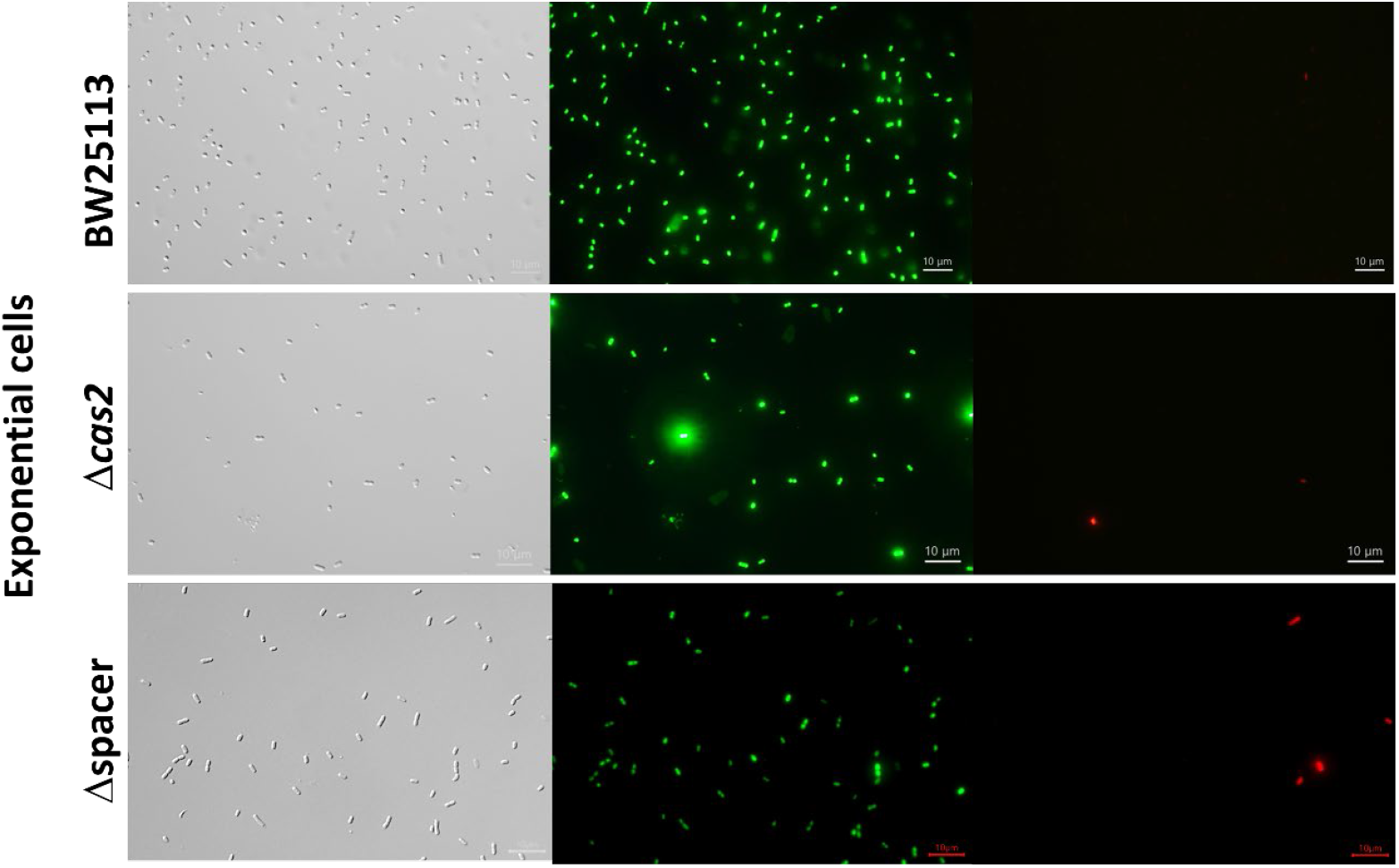
LIVE/DEAD staining of stationary (turbidity 2.0) and exponential (turbidity 0.8) cells shows the *cas2*, Δspacer (all 13 spacers removed), and mutations cause cell death. DF is dark field, SYTO9 is a membrane permeable stain for nucleic acids (green), and PI is propidium iodide, which is a membrane impermeable stain for the nucleic acids of dead cells (red). Tabulated cell numbers are shown in **Table S2**, and representative images from two independent cultures shown.

**Supplementary Figure 7.**
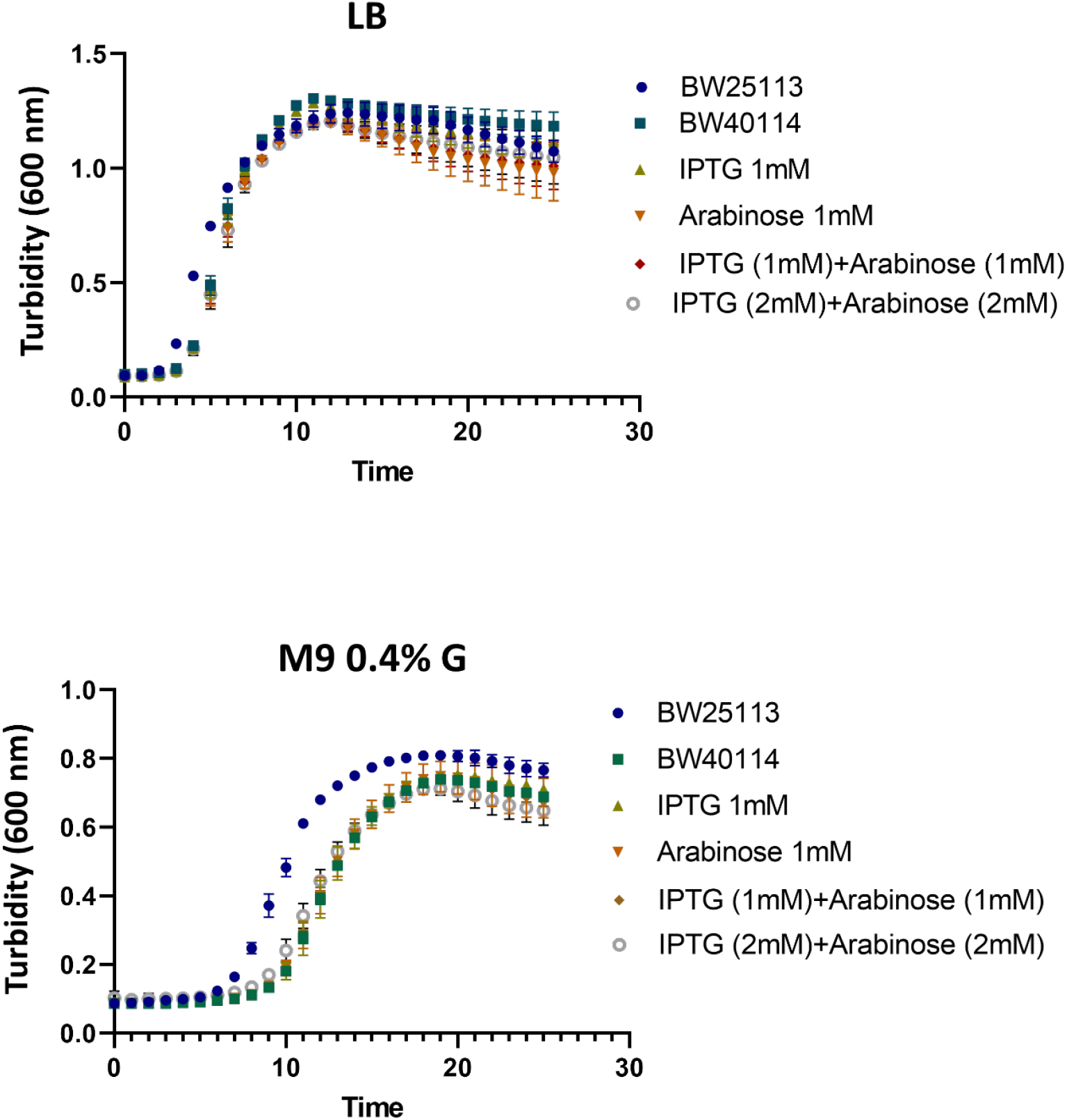
Growth curves upon inducing the full CRISPR-Cas system in *E. coli* BW40114 (via IPTG and arabinose) at 37°C in LB and M9 glucose media (turbidity at 600 nm shown). The growth is reduced due to the metabolic burden of producing the complete CRISPR-Cas system. Error bars indicate standard deviations from two independent cultures.

**Supplementary Figure 8.**
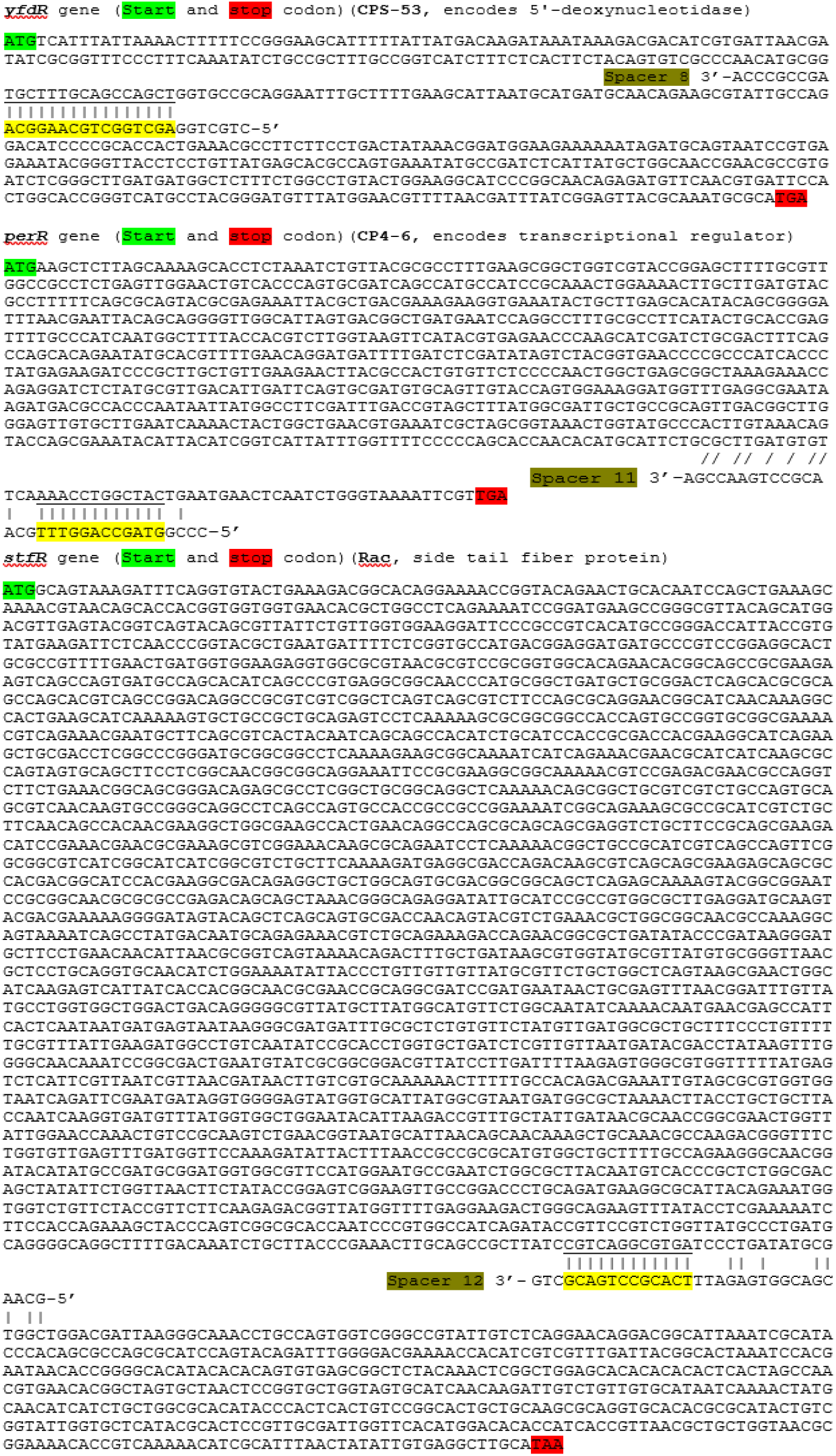
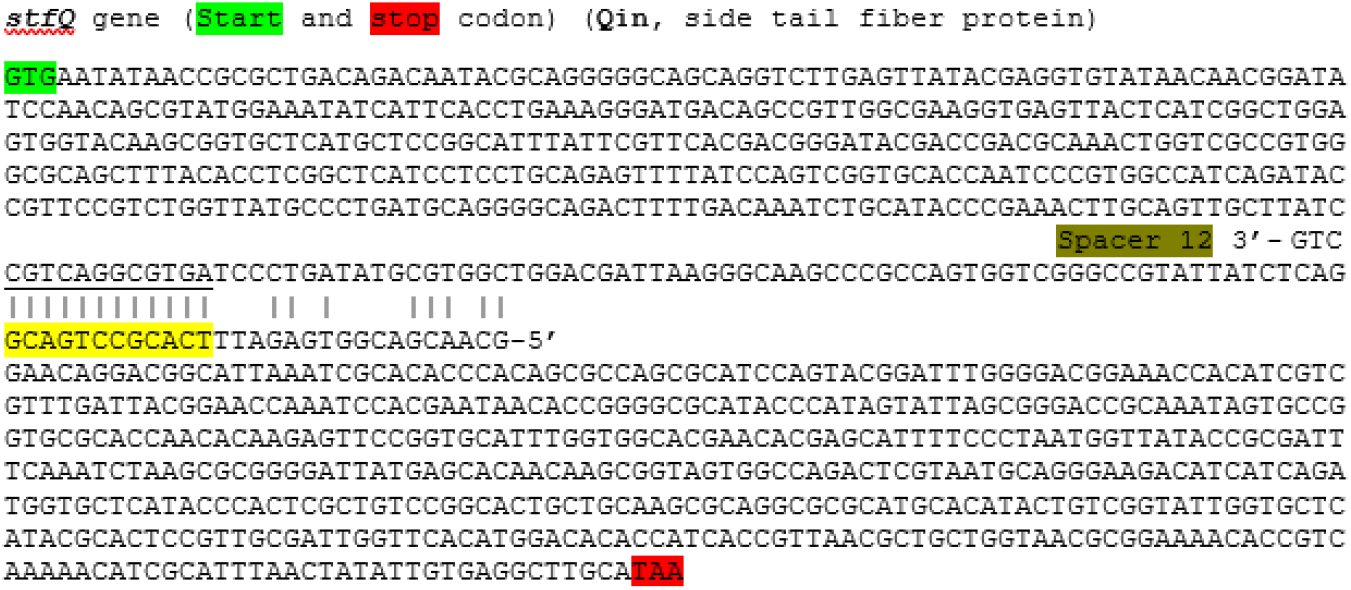
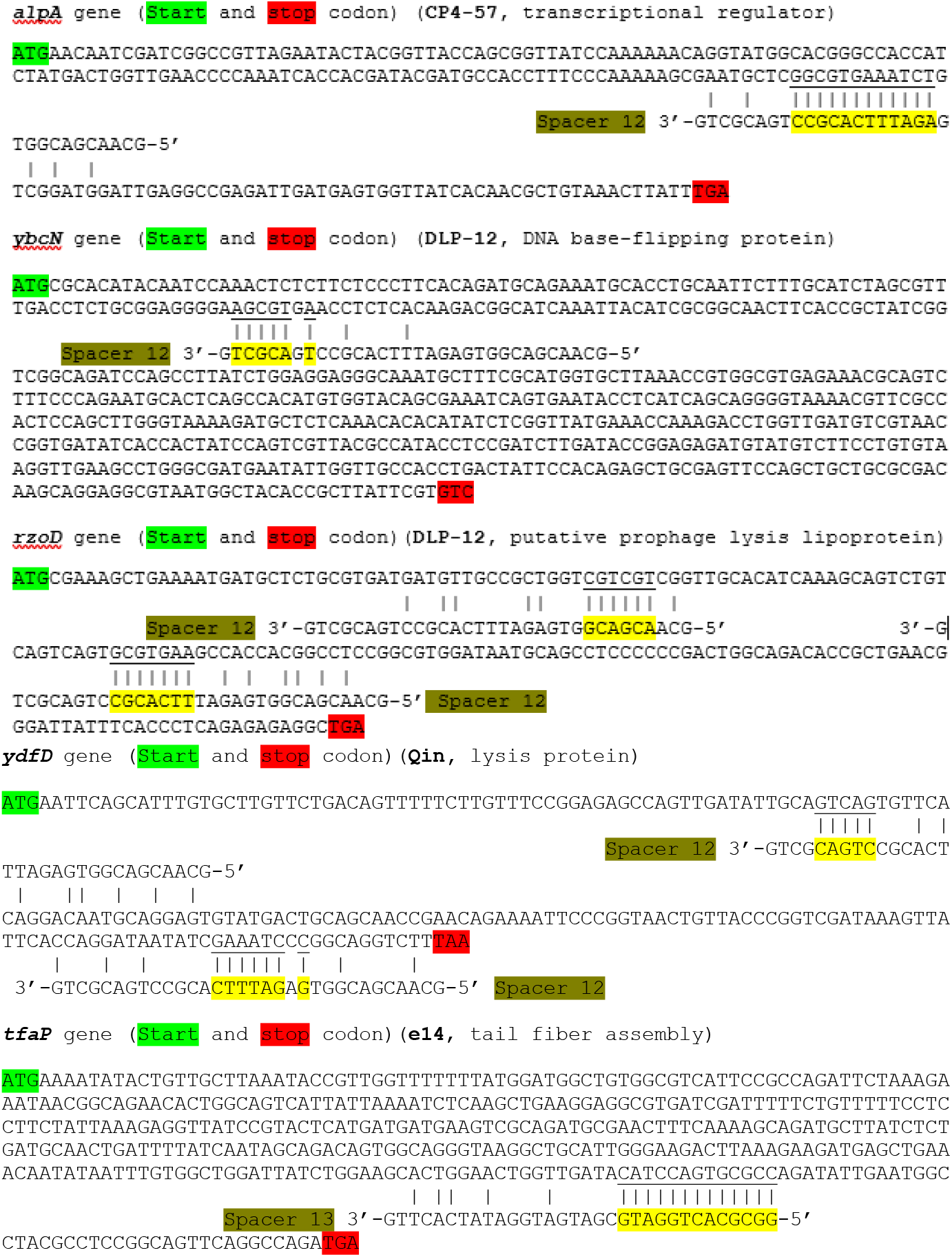
Predicted base-pairing interactions between each of the CRISPR spacers (i.e., spacers 8, 11, 12, and 13) and a protospacer within the cryptic phage mRNA (CPS-53, CP4-6, Rac, Qin, CP4-57, DLP-12, and e14). Note, some spacers match more than one region within a particular cryptic prophage. The complementary nt between the spacers and cryptic prophage protospacers is indicated by yellow highlight and vertical lines. Note that the spacer sequences are written 3’ to 5’ to aid an understanding of the putative RNAi matching.

**Supplementary Figure 9.**
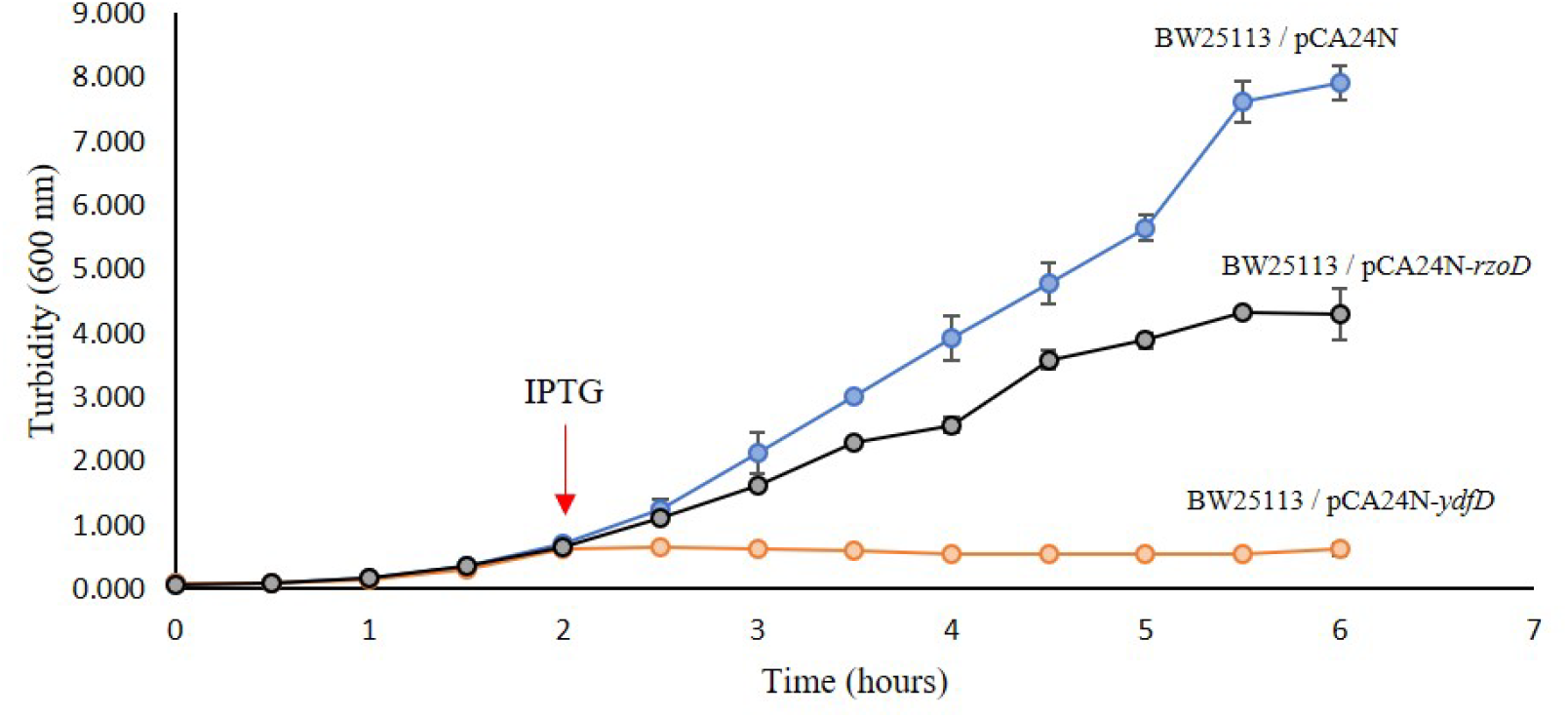
Toxicity of YdfD and RzoD. Growth curve at 37°C in LB (turbidity at 600 nm shown) of *E. coli* BW25113 harboring empty plasmid pCA24N (blue line), plasmid pCA24N encoding *ydfD* (orange line), and encoding plasmid pCA24N encoding for *rzoD*. Red arrow indicates addition of 1 mM of IPTG produce the toxins. Average of two independent cultures shown, and error bars indicate standard deviations.

**Supplementary Figure 10.**
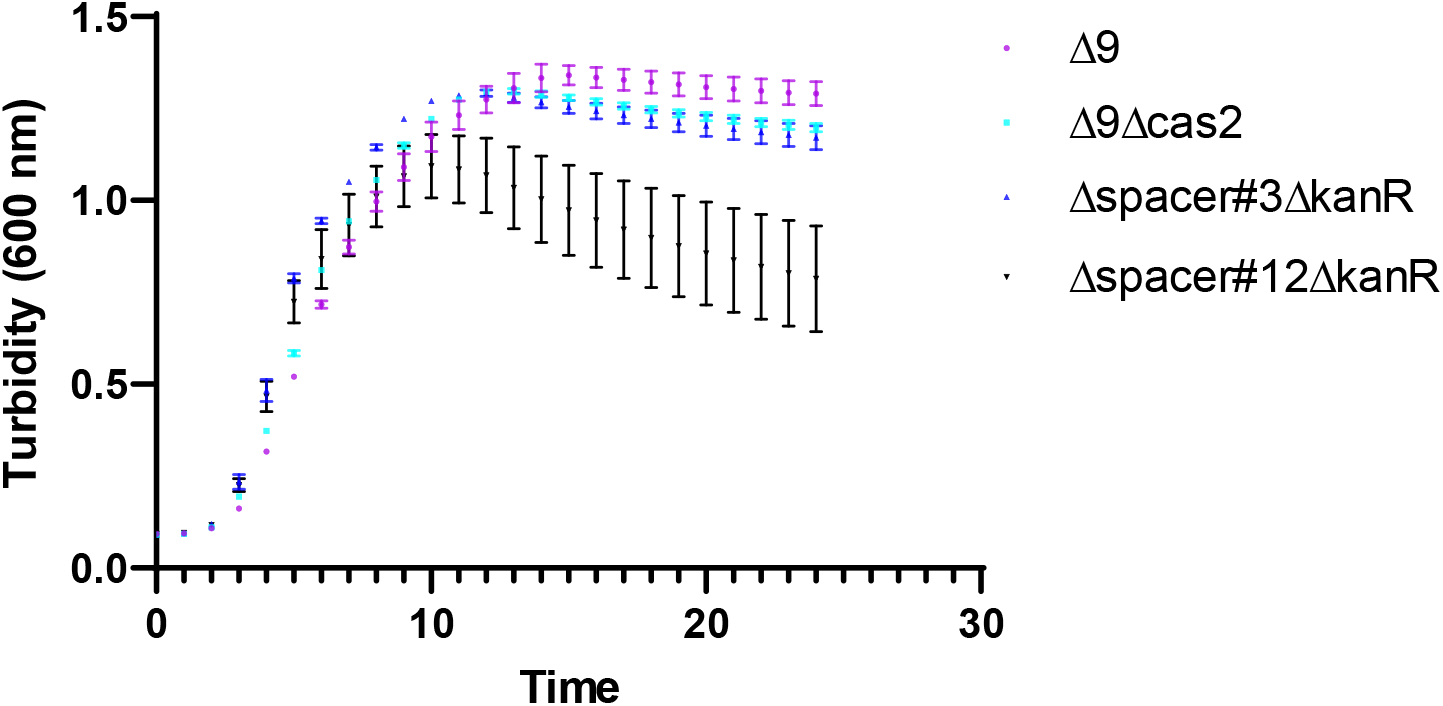
Deleting the DNA encoding spacer #12, which encodes crRNA targeting four separate regions in cryptic prophage mRNA (mRNA of Rac, Qin, CP4-57, and DLP-12), leads to lower cell yield. As seen **Table S2** and **Fig. S6**, this is associated with increased cell lysis. In contrast, deleting *cas2* in Δ9 and deleting spacer 3, which encodes crRNA that lacks matches in the mRNA of all 9 cryptic prophages has little effect on growth and cell lysis. Δ9 is BW25113 that lacks all cryptic prophages and Kan^R^ is kanamycin resistance. Growth (turbidity at 600 nm) at 37°C in LB shown, and error bars indicate standard deviations from two independent cultures.

**Supplementary Figure 11.**
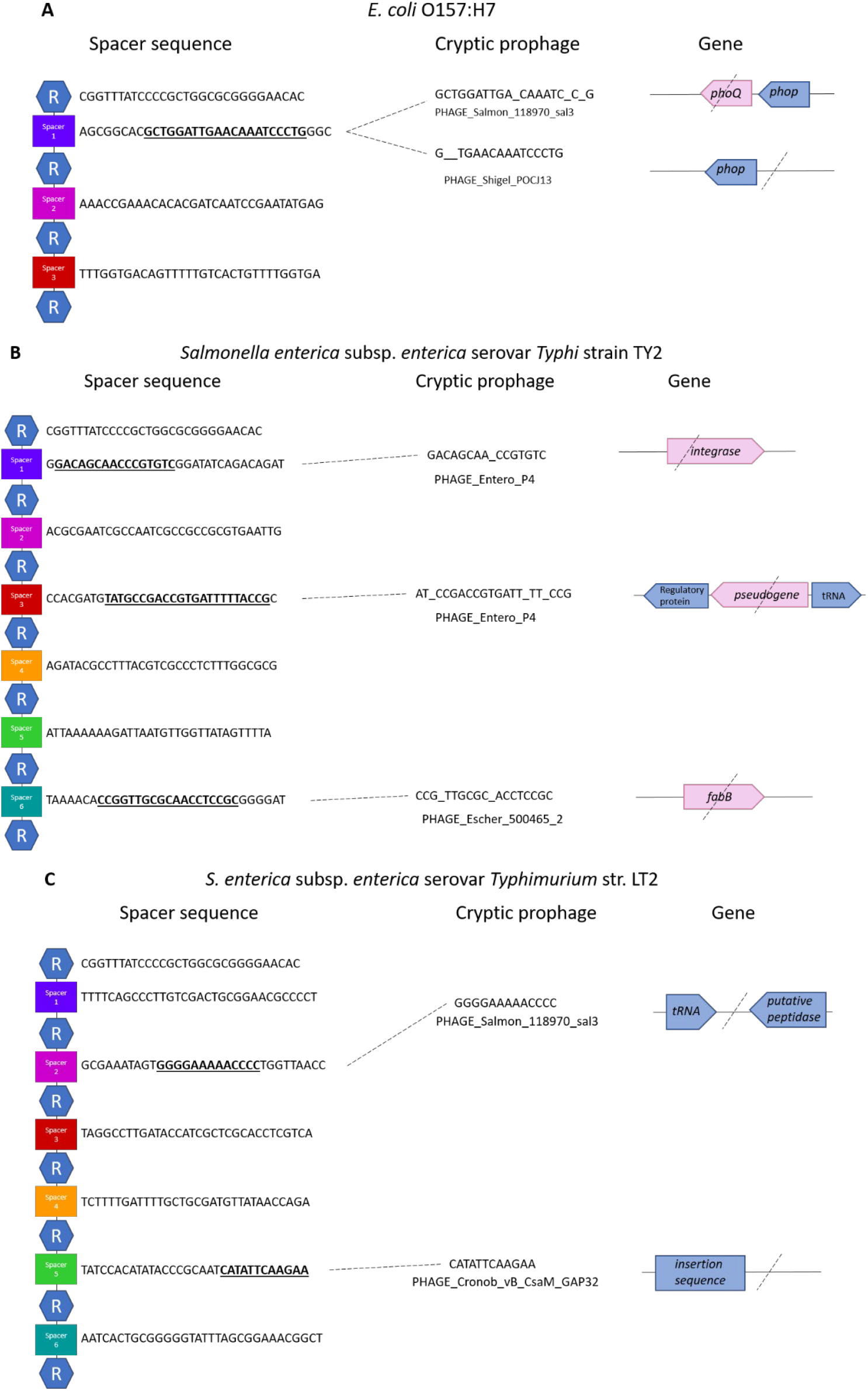
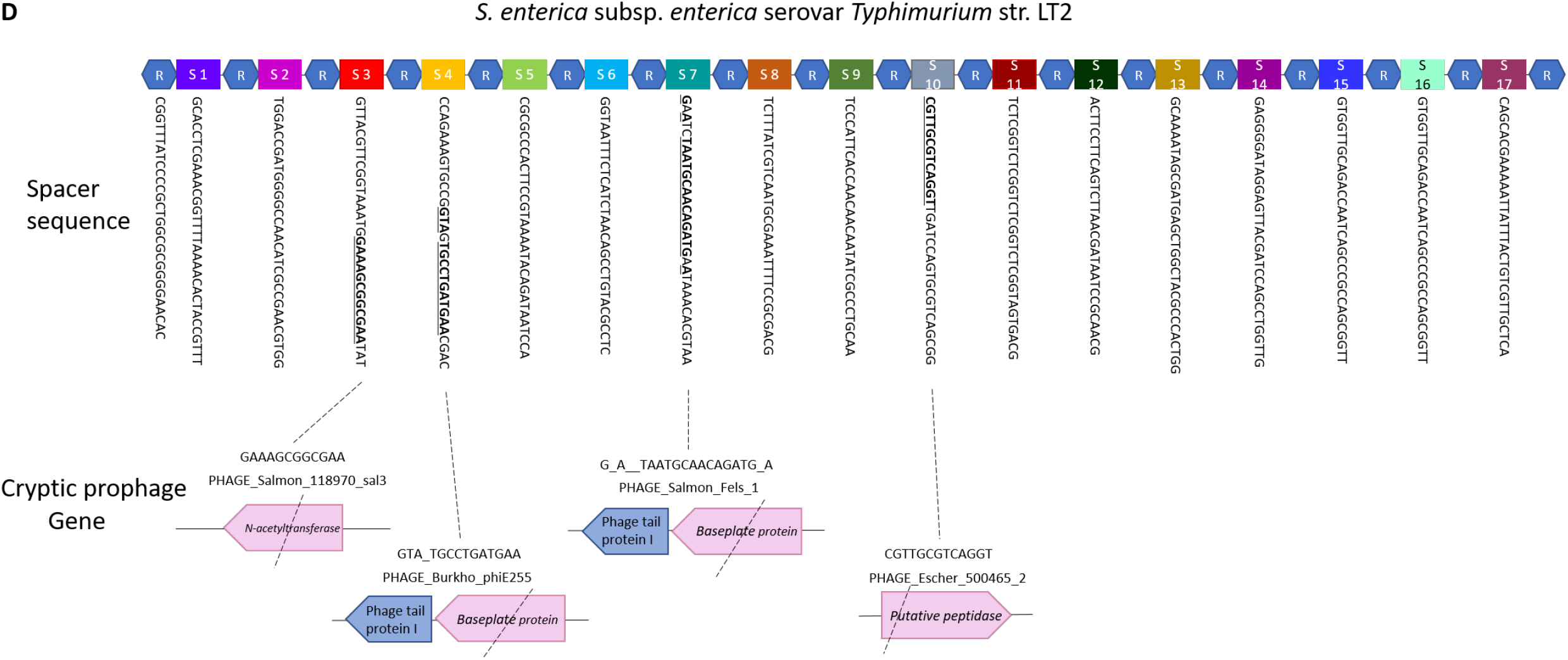
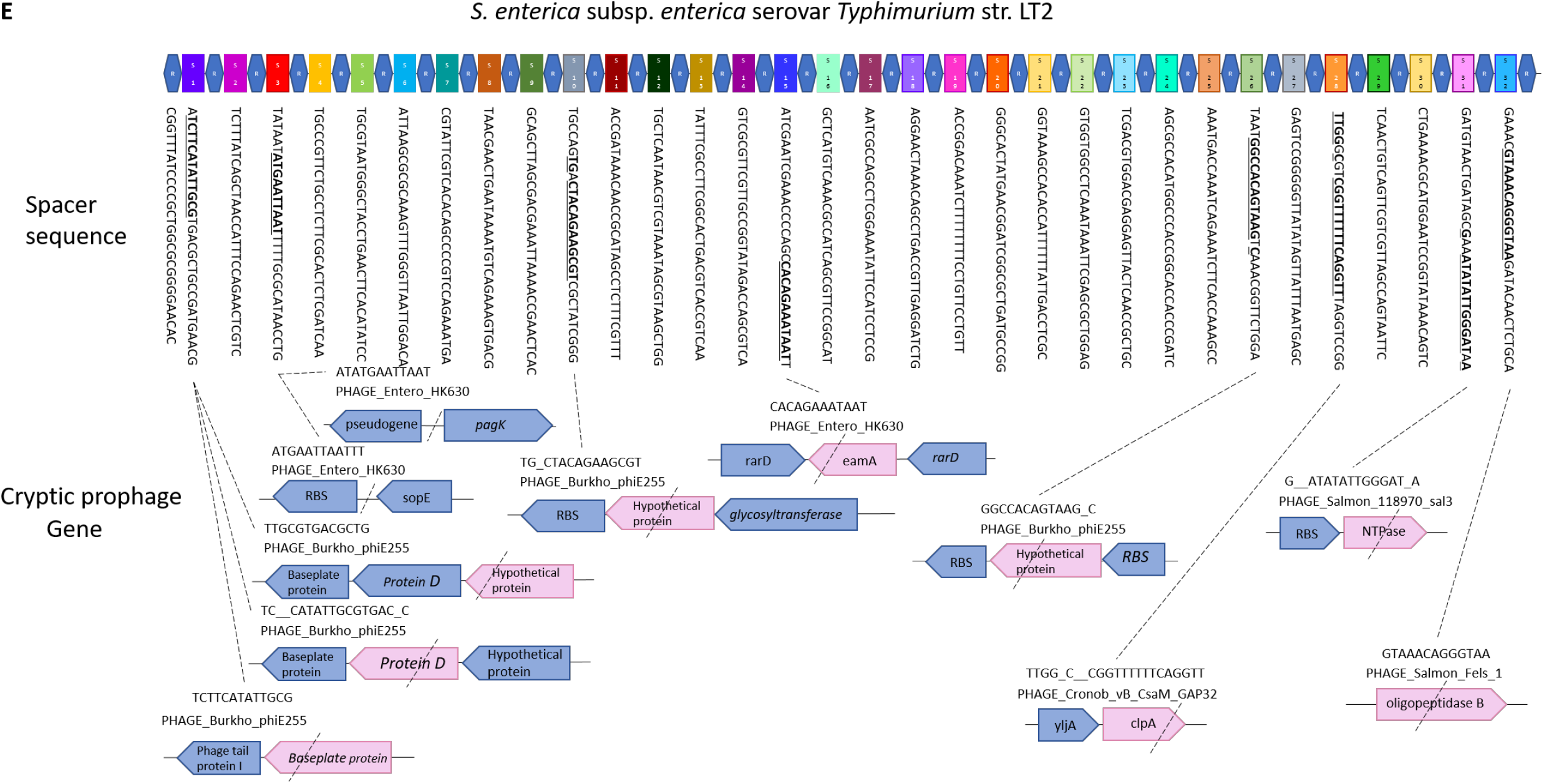
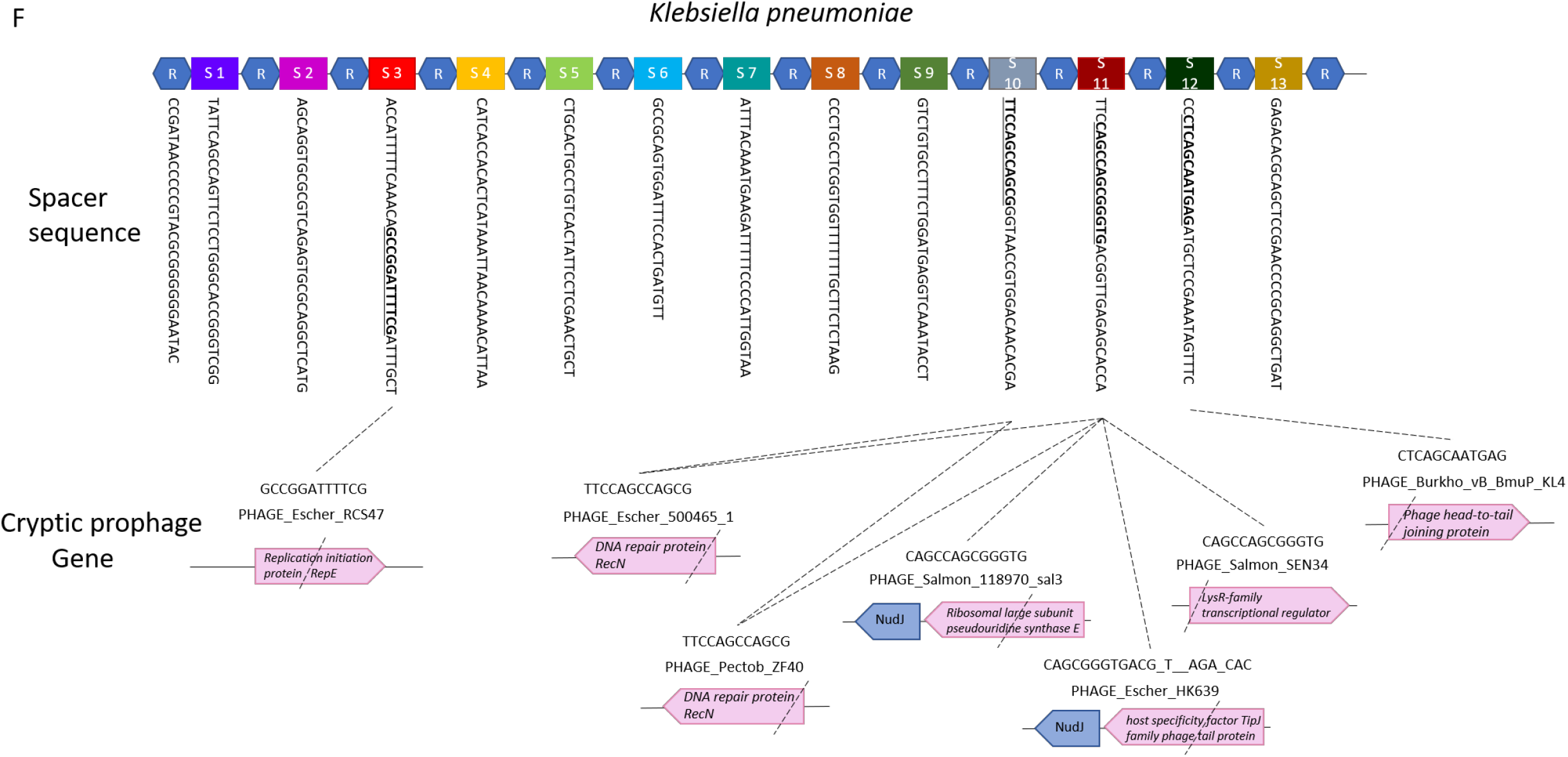
*E. coli* O157:H7 and *Salmonella* spp. CRISPR-Cas spacer sequences related to cryptic prophages. **(A)** Repeat (R, blue hexagon) and spacer (rectangle) sequences of each CRISPR-Cas system of *E. coli* O157:H7, *Salmonella* spp., and *K. pneumoniae* indicating matches with their cryptic prophages. (**A**) *E. coli* O157:H7, (**B**) *S. enterica* subsp. *enterica* serovar *Typhi* strain TY2, (**C, D, E**) three CRISPR-Cas systems of *S. enterica* subsp. *enterica* serovar *Typhimurium* str. LT2, and (**F**) *K. pneumoniae*. Pink highlight and dashed lines indicate the spacer positions relative to the cryptic prophage genes.

